# Preictal high-connectivity states in epilepsy: Evidence of intracranial EEG, interplay with the seizure onset zone and network modeling

**DOI:** 10.1101/2025.01.24.634461

**Authors:** Nicolas Medina, Manel Vila-Vidal, Ana Tost, Mariam Khawaja, Mar Carreño, Pedro Roldán, Jordi Rumià, María Centeno, Estefanía Conde, Antonio Donaire, Adrià Tauste Campo

## Abstract

**Objective.:** Epilepsy affects around 50 million people worldwide, and reliable pre-seizure biomarkers could significantly improve neuromodulation therapies for drug-resistant patients. Recent research using stereo-electroencephalography (sEEG) has revealed transient changes in network dynamics preceding seizures. In particular, our previous work showed that these alterations are driven by recurrent, short-lasting (0.6 s) high-connectivity network configurations—termed High-Connectivity States (HCS). Here, we aim to replicate and further characterize HCS as a biomarker in a multicentric patient cohort, assess its robustness across recording modalities and montages, explore its relationship with interpretable physiological variables, and examine its network-level association with seizure-onset zone (SOZ) dynamics.

**Approach.:** We analyzed long-term intracranial EEG (iEEG) recordings from 12 patients with sEEG and electrocorticography (ECoG). In two patients with extensive clinical information, we examined the interplay between HCS and SOZ dynamics. We also developed a low-dimensional stochastic network model to investigate mechanistic rationales of HCS emergence. Additionally, we compared HCS dynamics with gamma-band activity and heart rate, and tested robustness across different montage configurations.

**Main Results.:** In most patients, HCS probability reliably increased hours before seizure onset. In the two deeply characterized patients, this increase was specifically linked to an increased network centrality within the SOZ. The network model revealed that changes in HCS probability stem primarily from topological reconfigurations rather than changes in mean connectivity, underscoring the importance of dynamic interactions between epileptogenic and non-epileptogenic regions.

**Significance.:** These results support HCS probability as a promising biomarker for early seizure prediction and offer mechanistic insights into pre-seizure brain network dynamics.

## 1. INTRODUCTION

Epilepsy is one of the fourth most common neurological disorders worldwide, affecting approximately 0.5–1% of the global population (Sander, 2013). Among those affected, 30– 40% develop drug-resistant epilepsy, for whom surgical interventions remain the most effective treatment option (Sharma et al., 2015). A critical step in the treatment of these patients is the accurate identification of the seizure onset zone (SOZ) through continuous intracranial recordings such as stereo-electroencephalography (sEEG) or electrocorticography (ECoG) over up to three weeks. However, even with these invasive tools, real-time identification of pre-seizure periods still remains elusive, limiting the potential of adaptive neuromodulation therapies designed to anticipate and prevent seizures (Park et al., 2020).

Several studies have sought to characterize preictal brain dynamics by analyzing functional connectivity and network-based metrics derived from EEG (Lehnertz et al., 2023; Cousyn et al., 2023). These metrics include the use of clustering coefficients and centrality metrics to extract network metrics potentially linked to “seizure risk”. More recently, it has been shown that connectivity-based biomarkers derived from brief recordings can predict seizure likelihood with accuracy comparable to complex models (Khambhati et al., 2024). Nevertheless, many of these approaches show promise but face key challenges, including limited generalizability from small, homogeneous datasets; sensitivity of connectivity measures to non-stationarity; and a lack of physiological interpretability.

A key unresolved question is whether specific short-lasting network “states” can serve as robust and generalizable indicators of seizure risk. In prior work, Tauste Campo et al. (2018) demonstrated that brain network variability decreases prior to seizures, as reflected in a drop in centraility entropy. This effect is primarily driven by a higher likelihood of transient short-lasting (0.6s) high connectivity states (HCS) occurring during the hours preceding a spontaneous clinical seizure as compared to time-matched periods from previous days. This phenomenon reflects altered network dynamics in the pre-ictal phase and highlights the potential of HCS dynamics as a predictive biomarker of seizure risk extending beyond the activity of the SOZ alone.

Furthermore, the integration of physiological variables, such as heart rate or circadian rhythms, has shown promise in seizure forecasting (Vandecasteele et al., 2021). Notably, autonomic dysregulation, including seizure-induced tachycardia or bradycardia, has been observed in patients with temporal lobe epilepsy and could serve as a complementary pre-seizure biomarker (Billeci et al., 2018). However, few studies have combined the analysis of network states inferred from EEG with physiological markers, limiting our understanding of their joint predictive value for seizure forecasting (R. S. Fisher et al., 2014).

In the present study, we aim to extend and validate the HCS framework in three key directions. First, we perform general analysis across 12 patients using long-term intracranial EEG recordings —10 of them from an open database— to evaluate the reproducibility of preictal HCS dynamics on a larger, multicentric, heterogeneous cohort. Second, we focus in depth on two patients —one with SEEG and one with ECoG— with detailed clinical and recording data, to examine the relationship between HCS probability and interpretable internal and external variables such the relative activity of the SOZ with respect to remaining recorded sites or the heart-rate measurements, respectively. Third, we introduce a low-dimensional stochastic network model to investigate the topological mechanisms underlying the emergence of HCSs. This approach allows us to assess both the generalizability and physiological specificity of HCS-related network reorganization.

Taken together, our results provide new insights into the dynamics and correlates of preictal network reorganization. By doing so, this study contributes to the development of interpretable and multimodal seizure forecasting strategies.

## 2. METHODS

### Patients and recordings

This study utilized intracranial electroencephalography (iEEG) recordings obtained prior to the first recorded spontaneous clinical seizure from a multicentric cohort of 12 patients. Patient 1 (P1) underwent presurgical evaluation at the Epilepsy Unit of Hospital Clínic in Barcelona, Spain, and the clinical characteristics are summarized in Table 2. sEEG recordings were performed using a standard clinical intracranial EEG system (XLTEK, Natus Medical) with a sampling rate of 2048 Hz. Intracranial multichannel electrodes (Dixi Medical, Besançon, France; diameter: 0.8 mm; 5–18 contacts, contact length: 2 mm, inter-contact distance: 1.5 mm) were stereotactically implanted using frameless stereotaxy, neuronavigation assistance, and intraoperative CT-guided O-Arm imaging with the Vertek articulated passive arm. Electrode implantation decisions, target selection, and monitoring durations were determined based on clinical requirements. This patient provided informed consent for the use of their sEEG data in research. sEEG recordings included interictal periods spanning up to 2 days before the first marked seizure. Seizure onset and termination were defined electrographically through visual inspection by two epileptologists (AD, MK) and led to a surgical intervention (laser interstitial thermal therapy) with a very good post-surgical outcome (Engel I) after a follow-up of 2 years and a half (Table 2). We also analyzed ECoG data from a second patient (P2) with available clinical information through the public database IEEG.org. This patient was evaluated at the University of Pennsylvania Hospital and underwent implantation of intracranial electrodes for clinical monitoring. The electrode configuration included: a right parietal strip (RPT) with 4 contacts (RPT 1–4), a right interhemispheric strip (RIH) with 8 contacts (RIH 1–8), a right subtemporal strip (RST) with 4 contacts (RST 1–4), two 32-contact grids over the right temporal region (RG 1–64), and a left lateral strip (LL) with 8 contacts (LL 1–8). In both patients (P1 and P2), we had clinically validated and annotated seizure onset zone contacts.

Finally, we complemented this study with data from 10 additional patients from the publicly available SWEC-ETHZ iEEG long-term database (http://ieeg-swez.ethz.ch/) (Figure 3), produced through a joint initiative between the Sleep-Wake-Epilepsy-Center (SWEC) at the University Department of Neurology, Inselspital Bern, and the Integrated Systems Laboratory, ETH Zurich (ETHZ). This dataset consists of over 2,500 hours of continuous iEEG data including sEEG and ECoG recordings across 18 patients. To enable comparison of an 8-hour window prior to seizure onset with the same time window on the previous day, we included only patients with at least 32 hours of continuous recording prior to their first seizure. The selected patients, labeled as P3 to P12 for consistency, correspond to IDs 01, 02, 03, 11, 12, 14, 15, 16, 17, and 18 in ascending order from the original database. The number of electrodes varied across patients, ranging from 24 to 98 (Table 1).

**Table 1.**
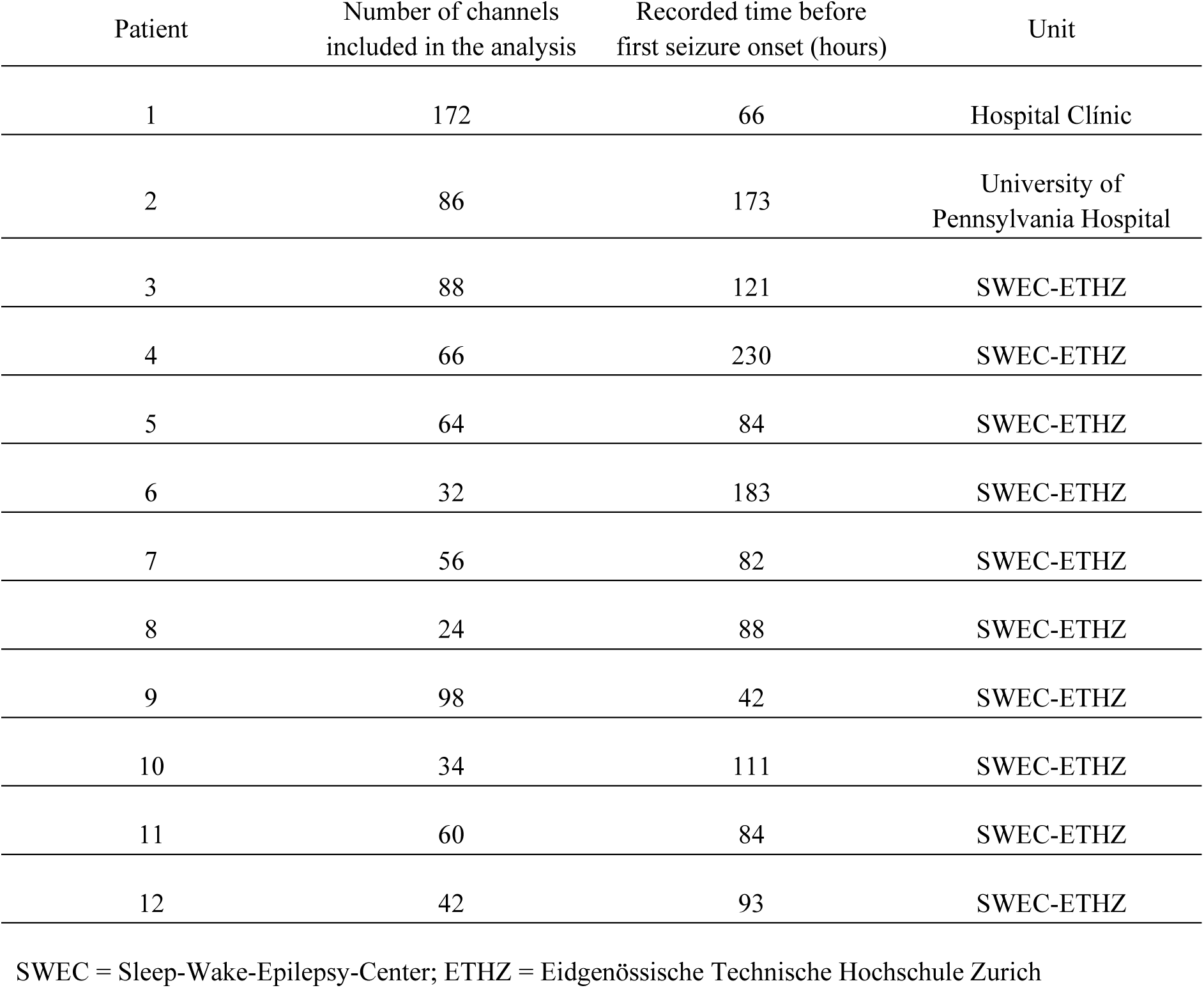
Common data of the 12 patients included in the study.

**Table 2.** Data of two patients with available clinical information.

| Patient | Gender<br>/Age | Epilepsy<br>onset<br>(years) | Suspected<br>epileptic focus | Hand<br>laterality | Implantation | Number of<br>electrodes<br>(right/left) | Number of<br>channels<br>included in the<br>analysis | Recorded time<br>before first<br>seizure onset<br>(hours) | Magnetic<br>Resonance | Surgery | Coregistered<br>ECG | Outcome<br>(2 y ½) |
| --- | --- | --- | --- | --- | --- | --- | --- | --- | --- | --- | --- | --- |
| 1 | M/30 | 18 | R lateral frontal | R | Bilateral | 3/14 | 172 | 66 | Normal | LITT | Yes | Engel I |
| 2 | M/26 | 15 | R neocortical<br>temporal | N/A | Bilateral | 48/8 | 86 | 173 | N/A | N/A | Yes | N/A |

ECG = electrocardiogram; R = right; LITT = laser interstitial thermal therapy; N/A= not available information.

### Data preprocessing

Data were obtained from three databases: Hospital Clínic, IEEG.ORG, and SWEC-ETHZ iEEG. For each case, recordings spanning up to 32 hours prior to seizure onset were included in the analysis. Channels exhibiting artifacts were visually identified and excluded. To improve computational efficiency, all data were uniformly downsampled to 512 Hz, irrespective of the original sampling frequency. To minimize the influence of slow drifts and baseline shifts, each EEG segment was linearly detrended prior to correlation analysis. This procedure involved fitting and subtracting a linear trend from each signal, effectively removing low-frequency components while preserving transient fluctuations relevant to functional connectivity estimation. This is particularly important in signal processing and time series analysis, where trends can obscure meaningful patterns or introduce artifacts. Consequently, depending on the specific analysis, a band-pass Butterworth filter (1–150 Hz, 53 dB stopband attenuation, maximal ripples in passband 2%) was applied. Additionally, a notch filter at 50 Hz and its harmonics was used to eliminate power line interference.

### General procedure

The general objective was to evaluate the presence and significance of transient high-connectivity states (HCSs) in the hours preceding seizure onset using intracranial EEG. We first assessed this phenomenon in a cohort of 12 patients, including both stereo-EEG and ECoG recordings, by comparing network variability (centrality entropy) and HCS probability on the seizure data with the same time window on the previous day. This analysis aimed to replicate and extend the findings of our previous study (Tauste Campo et al., 2018), where a sustained and significant preictal drop in entropy was observed, primarily due to an increased likelihood of HCSs.

After establishing cohort-level effects, we conducted a more detailed analysis in two well-characterized patients –P1, with sEEG; P2, with ECoG recordings– for whom long-term heart rate was available. In these cases, we explored the correlation between HCS probability with other physiological and network-based variables, including signal power, heart rate, mean connectivity, and mean node centrality.

### Power analysis

To compute normalized mean power, a baseline segment was first defined as the initial 5 minutes of recording on the first day to obtain the baseline power mean and standard deviation (𝑃_𝑏𝑎𝑠𝑒𝑙𝑖𝑛𝑒_). After preprocessing the data, we estimated the power by integrating the power spectral density in the frequency band of interest according to previous studies (Vila-Vidal et al., 2020). Finally, we normalized the power values with respect to the baseline power (𝑃_𝑏𝑎𝑠𝑒𝑙𝑖𝑛𝑒_) considering all channels. Then, we calculated the Z-score values (𝑍) using the mean and standard deviation of the baseline power (See Fig. S1 for a scheme illustration).

### Functional connectivity analysis

We began by estimating functional connectivity from intracranial EEG signals. A widely used approach for this is Pearson correlation, which captures linear dependencies between time series. Pearson correlation measures the strength and direction of the linear relationship between two signals by calculating the correlation coefficient, which ranges from -1 to 1, indicating perfect negative to perfect positive linear correlation, respectively (Pereda et al., 2005). This approach has been extensively applied in neuroimaging to construct functional connectivity networks by correlating the activity between brain regions (Friston, 2011). Its simplicity enables easy implementation and interpretation, although it may miss nonlinear interactions (Wendling et al., 2009).

Let 𝑥 and 𝑦 be 2 𝑁 -length time series representing 2 recorded signals and let 𝑥̅ and 𝑦̅ be their respective sample means. Pearson correlation is computed as follows:

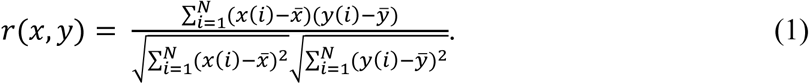

For each patient and each consecutive 0.6-second window, we computed the absolute value of the correlation coefficient across all pairs of electrode channels, yielding a single MxM functional connectivity (FC) matrix for each 0.6 s, where M is the total number of electrode channels. These windowed correlation matrices (sampled at an approximate frequency of 1.67 Hz) served as the basis for subsequent network-level analyses.

### Network states analysis

To capture time-varying network configurations, we analyzed network states using eigenvector centrality, a graph-theoretical measure of node importance. This approach follows the methodology described in Tauste Campo et al. (2018). For a given graph 𝐺 = (𝑉, 𝐸), let 𝐴 = (𝑎_𝑣_, 𝑡) be its weighted adjacency matrix. The relative centrality score 𝑥_𝑣_ of vertex 𝑣𝜖𝑉 can be defined as:

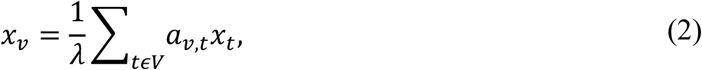

which can be rearranged in a matrix form as 𝜆𝑥 = 𝐴𝑥 . According to the Perron-Frobenius theorem, the largest eigenvalue yields a valid centrality measure, provided that all entries in 𝑥 are non-negative (Newman, 2010). The centrality measure is determined by the eigenvector corresponding to the largest eigenvalue of the absolute-valued connectivity matrix. The centrality value for the 𝑖 -th channel is the 𝑖 -th component of this eigenvector, with 𝑖 ranging from 1 to the number of recording sites in a patient. This measure is self-referential: nodes attain high centrality if they connect to other high-centrality nodes (Rubinov & Sporns, 2010), offering insight into each channel’s relative importance within the network.

By computing eigenvector centrality for each 0.6-second FC matrix, we generated an independent centrality time series for all recorded channels across the entire session (one centrality value every 0.6 seconds). Each FC matrix represents a distinct network state (Allen et al., 2014), and changes in the matrix induce a rotation of the eigenvector, dynamically updating the channels’ relative importance within the network (Tauste Campo et al., 2018). To quantify how these centrality patterns evolve over time, we estimated the multivariate Gaussian entropy of the centrality time series within consecutive, non-overlapping 180-second windows (300 centrality samples), as a second-moment approximation of network variability. The entropy is defined as:

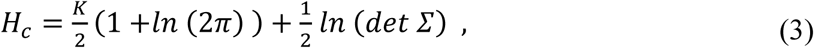

where 𝐾 is the number of recording sites, and 𝛴 is the covariance matrix of the centrality time series within the estimation window. In line with our previous findings, a significant decrease in centrality entropy prior to seizure onset is attributed to the increased occurrence of high-connectivity states (HCS).

To identify spontaneous network states over long interictal periods (>1 day), we applied k-means clustering to the centrality vectors. While k-means is computationally efficient and simple, it requires predefining the number of clusters 𝑘, and the algorithm’s convergence may depend on the initial placement of centroids (Bizopoulos et al., 2013). In this study, the optimal number of clusters 𝑘 was determined individually for each patient using the elbow method (Horvat et al., 2021). Note that network states were defined in a patient-specific manner to account for patient-specific implantation schemes. As a result of this analysis, we obtained a time series of 0.6-second resolution —yielding 6000 consecutive FC matrices per hour—, each classified into one network state over time.

### Heart rate analysis (only patients P1 and P2)

The heart rate is defined as the number of heart beats per minute. The duration of each beat (𝑡_𝑏𝑒𝑎𝑡_) is usually estimated from the time interval between consecutive 𝑅 -peaks. Heart rate analysis is a crucial component in the assessment of physiological states and the detection of potential abnormalities (Li et al., 2023). Preprocessing an ECG signal is crucial for accurate heart rate calculation, as it removes noise and artifacts that can obscure the detection of 𝑅 - peaks, which are pivotal for determining heart rate. The process typically involves filtering to remove baseline wander and power line interference, using high-pass, notch, and band-pass filters (Clifford, 2006). First, we down-sampled the raw ECG data from 2048 to 512 Hz. We mitigated the Baseline wander, caused by respiration and patient movement, with high pass Butterworth filtering around 0.5 Hz (53 dB stopband attenuation, maximal ripples in passband 2%). Then, we applied a notch filter at 50 Hz to remove the power line interference. Afterward, we use a bandpass Butterworth filter (53 dB stopband attenuation, maximal ripples in passband 2%) from 0.5 to 40 Hz to refine the signal by eliminating irrelevant high-frequency noise. To enhance the QRS complex and facilitate robust R-peak detection, additional processing steps, including differentiation, squaring, and moving window integration, were applied, following the Pan and Tompkins method (Pan & Tompkins, 1985). The heart rate (𝐻𝑅) was estimated as 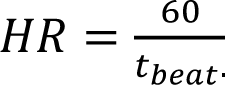. where 𝑡_*beat*_ is the interval between consecutive R-peaks (See Fig. S1 for a scheme illustration).

### Statistical analysis

To evaluate significant and sustained differences in time-varying measures such as mean power, mean connectivity, and centrality entropy, we employed a nonparametric cluster-based statistical test (Maris & Oostenveld, 2007). This test quantifies time-matched changes between variables recorded on the seizure day and the previous day. Specifically, we partitioned the continuous time series into non-overlapping segments of 20 minutes, corresponding to 20 consecutive biomarker samples. For each segment pair, we performed a Wilcoxon test (1,000 random permutations) to determine statistical significance. Adjacent segments meeting a significance threshold of 𝑃 < *0*.*05* over a minimum duration of 3 segments (1 hour) were grouped into clusters, and the mean significance within each cluster was considered the primary test statistic. The null distribution was obtained by calculating the test statistic for all possible reorganizations of the data groups (exact test).

To investigate the relationship between HCS probability and centrality entropy reduction (Fig. 2), we applied linear regression analysis and calculated the coefficient of determination (𝑅*^2^*). Additionally, mean connectivity values across electrode pairs were Fisher-transformed to stabilize variance and normalize the correlation coefficients (R. A. Fisher, 1915).

### Low-dimensional stochastic network model

We proposed a simplified stochastic network model to reproduce the time-dependent emergence of high-connectivity states (HCS) in the preictal period. The goal of this network model is to isolate the role of network topology in driving changes in HCS probability, independently of changes in average connectivity. The model constraints mean functional connectivity and mean centrality values across non-overlapping time windows during the 4 hours preceding seizure onset, both on the seizure day and the corresponding period of the previous day. More specifically, the model works by generating continuous signals whose sequential correlation matrices (normalized covariance matrices) are adjusted to meet predefined ranges for mean connectivity and centrality. To achieve a specific mean connectivity, the correlation matrix coefficients are iteratively adjusted until the desired mean connectivity value falls within a specified range. Next, to control mean centrality, the positions of the matrix coefficients are redistributed while keeping the mean connectivity constant. By altering the node positions rather than their values, mean centrality can be modulated independently of mean connectivity.

The model divides the preictal period into two stages: (1) From 2 to 4 hours before seizure onset, mean connectivity and mean centrality remain within the same ranges for both days, 0.2±0.01 for mean connectivity and 0.4±0.009 for mean centrality. (2) In the final 2 hours (from 0 to 2 hours before the seizure), the mean connectivity remains fixed at 0.25±0.01 for both days, while mean centrality increases only on the seizure day to 0.55±0.01.

From the covariance matrices generated by this model, eigenvector centrality values were extracted, enabling the identification of network states for both days following the same procedure applied to the real iEEG recordings. Subsequently, we calculated the time-varying probability of HCS. To ensure robust outcomes, the model was run 40 times, and results were averaged across all simulations.

## 3. RESULTS

### Pre-seizure HCS probability alterations: detailed analyses in patients P1 and P2

To investigate the pre-seizure specific emergence of high-connectivity states (HCS) in the new cohort of patients, we adopted the methodology from (Tauste Campo et al., 2018) (Fig. 1(a)). In addition, Figure 1(b) shows time-varying calculations of mean power, connectivity, centrality, and centrality entropy for an exemplary patient (P1) on the seizure day and the preceding day, focusing on the four hours leading up to seizure onset. In (Tauste Campo et al., 2018), centrality entropy was shown to decline prior to seizure onset, correlating with an increase in HCS occurrence during this period.

**Fig. 1.**
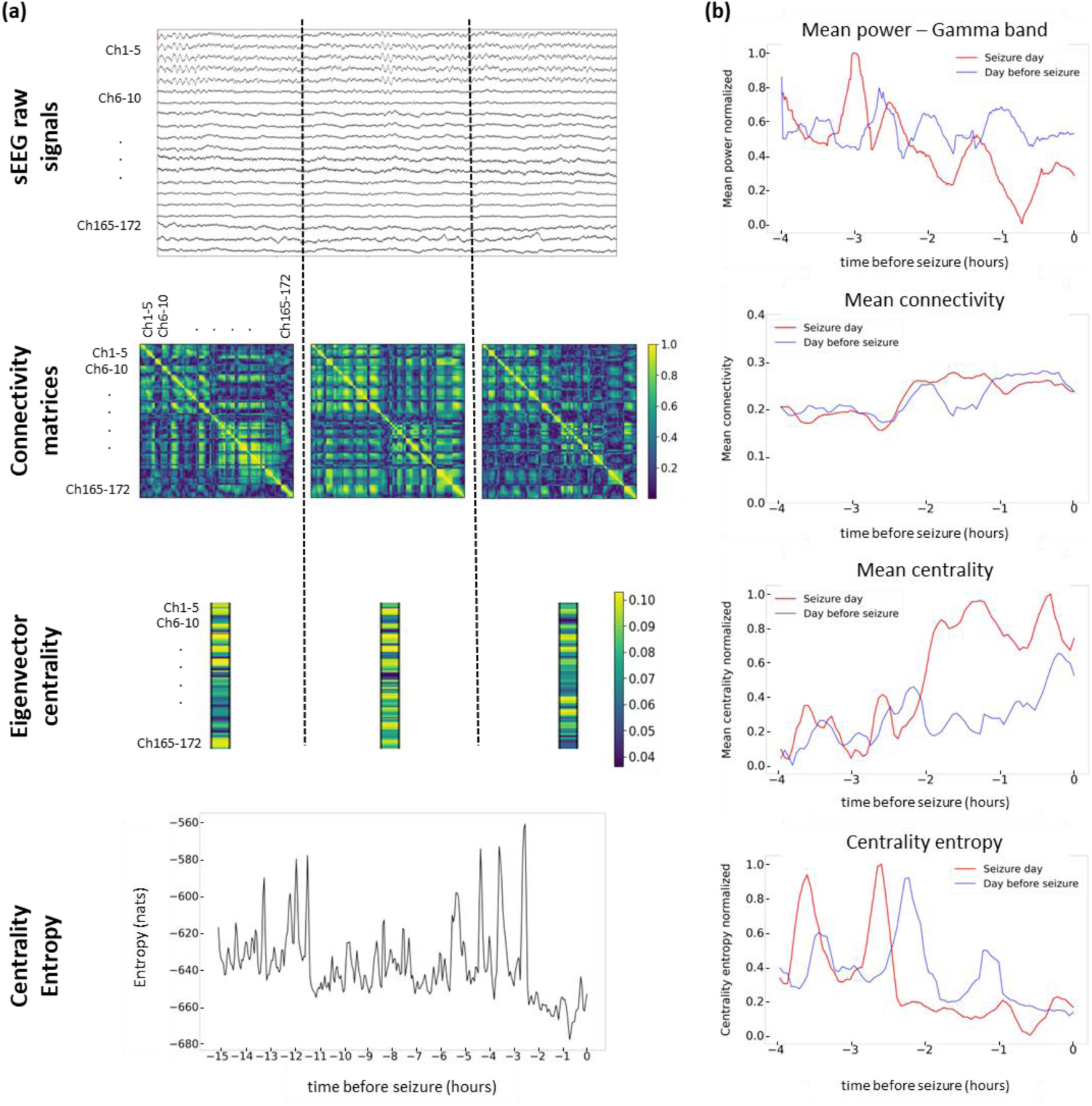
Time-matched continuous sEEG network dynamics analysis across subsequent days. The procedure used in the previous study (Tauste Campo et al., 2018) to analyze the dynamics of the sEEG network during long continuous recordings is described step-by-step. 1(a) Top: Each sEEG signal is divided into consecutive and non-overlapping time windows of 0.6 seconds duration. The channels are grouped according to their corresponding electrodes. Top middle: Functional connectivity matrices are computed using the absolute value of the Pearson’s correlation in each time window. Bottom middle: Eigenvector centrality is then calculated from each functional connectivity matrix to characterize transient network states. This approach transforms the 𝑁𝑥𝑁 order matrix into an 𝑁 -component vector that characterizes the network, where 𝑁 is the number of channels. Bottom: The centrality entropy values are computed for each period of interest by sequentially estimating the multivariate Gaussian entropy of the eigenvectors in consecutive and non-overlapping time windows of 180 seconds (300 samples). 1(b) For an exemplary patient (P1) with 172 electrode channels, time-varying channel-averaged normalized outputs of each procedure step during the seizure day (in red) and the day before (in blue). Top: normalized mean power. Top middle, mean connectivity. Bottom middle: normalized mean centrality. Bottom: normalized centrality entropy. The centrality entropy values are also shown with the sampling rate of 1 sample every 3 minutes. The 𝑥-axis represents the time prior to the seizure, with the value 0 indicating the instant of seizure onset (reference). Ch, channel.

Figure 2(a) illustrates the centrality entropy for P1 on the seizure day and the preceding day, where a significant decrease in centrality entropy emerged approximately 2.5 hours before seizure onset. To define HCS, we employed the k-means clustering algorithm on centrality eigenvectors, ranking states in descending order of mean connectivity estimated via the Fisher transform (R. A. Fisher, 1915). Twelve distinct states were identified using K-mean clustering and the elbow method (Horvat et al., 2021) and ordered according to the mean connectivity value of their corresponding centroids. Thus, State 1 corresponded to the state with highest mean connectivity (HCS) while State 12 corresponded to the state with lowest mean connectivity value (Fig. 2(b)). Next, we determined which state best explained the observed entropy decline using a linear regression model. Here, the dependent variable was the centrality entropy difference across days while the independent variable was the probability difference of each state. Figure 2(c) shows the coefficient of determination (𝑅^2^), revealing that the HCS (State 1) was the primary driver of the entropy drop. We then analyzed the HCS probability during the pre-seizure entropy decline window (0.5–2.5 hours before seizure onset), observing a significant increase in HCS probability on the seizure day compared to the previous day (randomization test, 𝑃 < 0.01). Figure 2(e) summarizes the time-varying dynamics of HCS probability over the 8 hours preceding seizure onset, highlighting a significant sustained increase in HCS probability as seizure onset approached, compared to the day before.

**Fig. 2.**
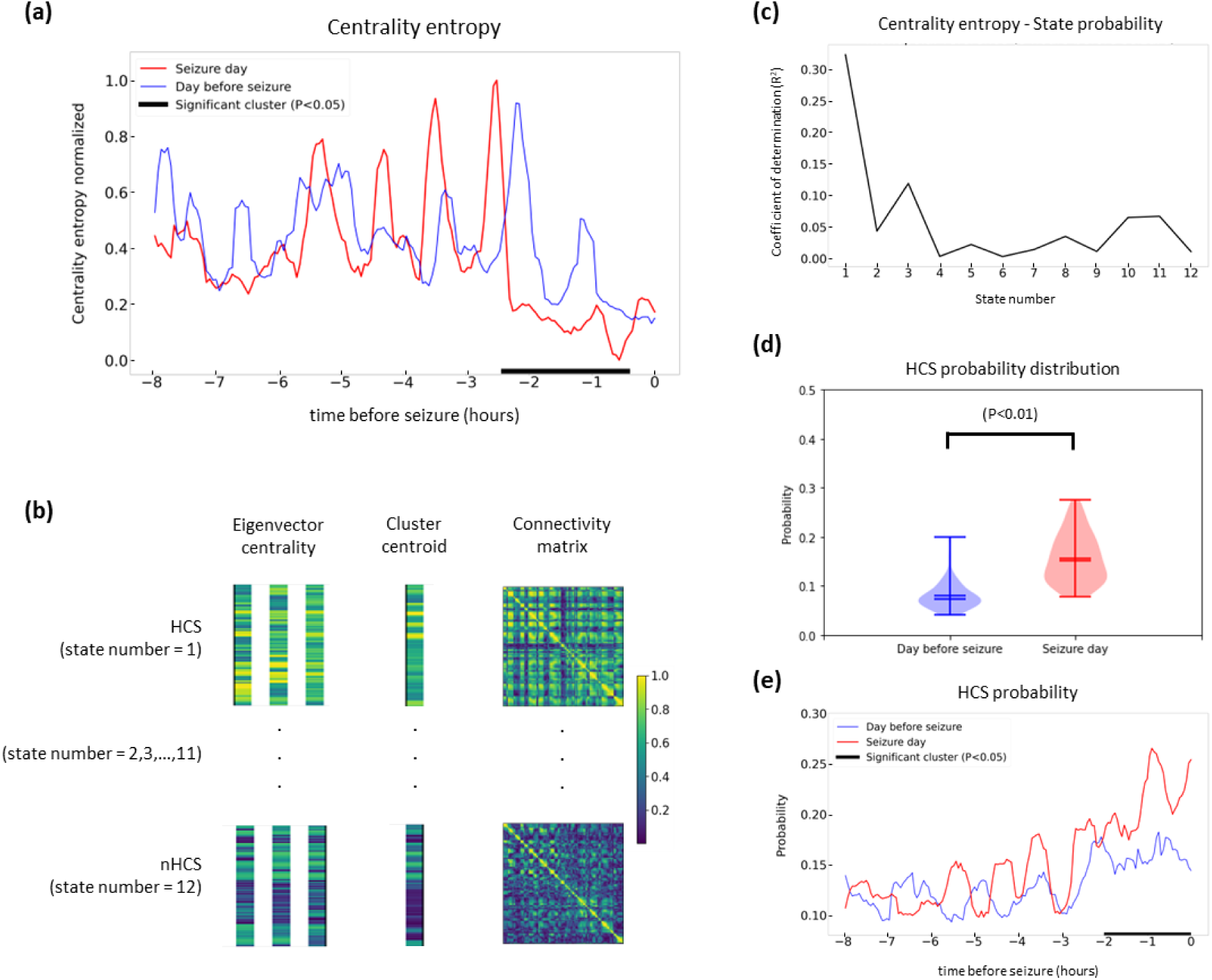
Increased probability of high connectivity states (HCS) explains changes in pre-seizure sEEG network variability (Patient 1). 2(a) Centrality entropy was calculated for the day of the seizure (red) and the previous day (blue). The 𝑥 -axis represents the time prior to the seizure in hours, with the value 0 marking the instant of seizure onset (reference). In this patient, a significant decrease in centrality entropy was observed from 2.5 to 0.5 hours before seizure onset (𝑃 < 0.01). 2(b) Definition of HCS and nHCS: Considering the entire data record, twelve states were defined and grouped in descending order according to their level of connectivity, with the first state corresponding to HCS and the last to the lowest connectivity. To cluster the centrality eigenvectors, an unsupervised 𝐾 -means clustering algorithm was employed. Once the centrality centroids were defined for each 600 ms time window, they were ordered in decreasing order based on their mean connectivity. 2(c) The coefficient of determination (𝑅^2^) was calculated for a linear regression model where the dependent variable was the centrality entropy, and the independent variable was the state probability. The state that best explains the entropy drop is HCS (State 1), with an 𝑅^2^ value above 0.3. 2(d) The distribution of HCS probability for both days during the critical period (from 0.5 to 2.5 hours before seizure onset) is shown. A significant increase in HCS probability (𝑃 < 0.01) is observed on the day of the seizure. 2(e) Finally, the dynamics of HCS probability over the 8 hours pre-ictal stage are depicted, illustrating a gradual increase in this parameter as the seizure onset approaches. The temporal window of significance occurs 2 hours before seizure onset (𝑃 < 0.05) which overlaps with the time window of the significant decrease of centrality entropy. In 1(a) and 1(e), the cluster-based permutation test was used to determine significance.

Next, we applied the above analysis in Patient 2 (P2) monitored with ECoG recordings (strips and grids) and most clinical information available. As illustrated by Figure S2, this independent case similarly demonstrated progressive decreases in centrality entropy (Fig. S2a) and increases in HCS probability (Fig. S2c) with these changes emerging 8 hours before the seizure onset in a linked fashion (Fig. S2b). To further validate the above findings, we reproduced the analyses using the publicly available SWEC-ETHZ iEEG database (18 patients) including patients with sEEG and ECoG recordings but with limited clinical information due to strict internal anonymization policy. From this database, 10 patients (P3–P12) met the inclusion criterion of having at least two days of recordings prior to the first monitored seizure. In 8 patients (P3–P7, P9–P10, P12), a pre-seizure decline in centrality entropy was observed (Fig. 3(a)), replicating the results seen in P1 and P2 (Fig. 2(a) and Fig. S2a). Over all patients, HCS (State 1) consistently explained the entropy drop during the seizure day (n=12, Mann-Whitney U paired Test, 𝑃 < *0*.*01*, Fig. 3(b)) significantly better than the remaining states on average. Finally, the time-varying HCS probability revealed a patient-specific increase aligned with the entropy decrease on the seizure day (except for P7 and P10), supporting our hypothesis (Fig. 3(c)).

**Fig. 3.**
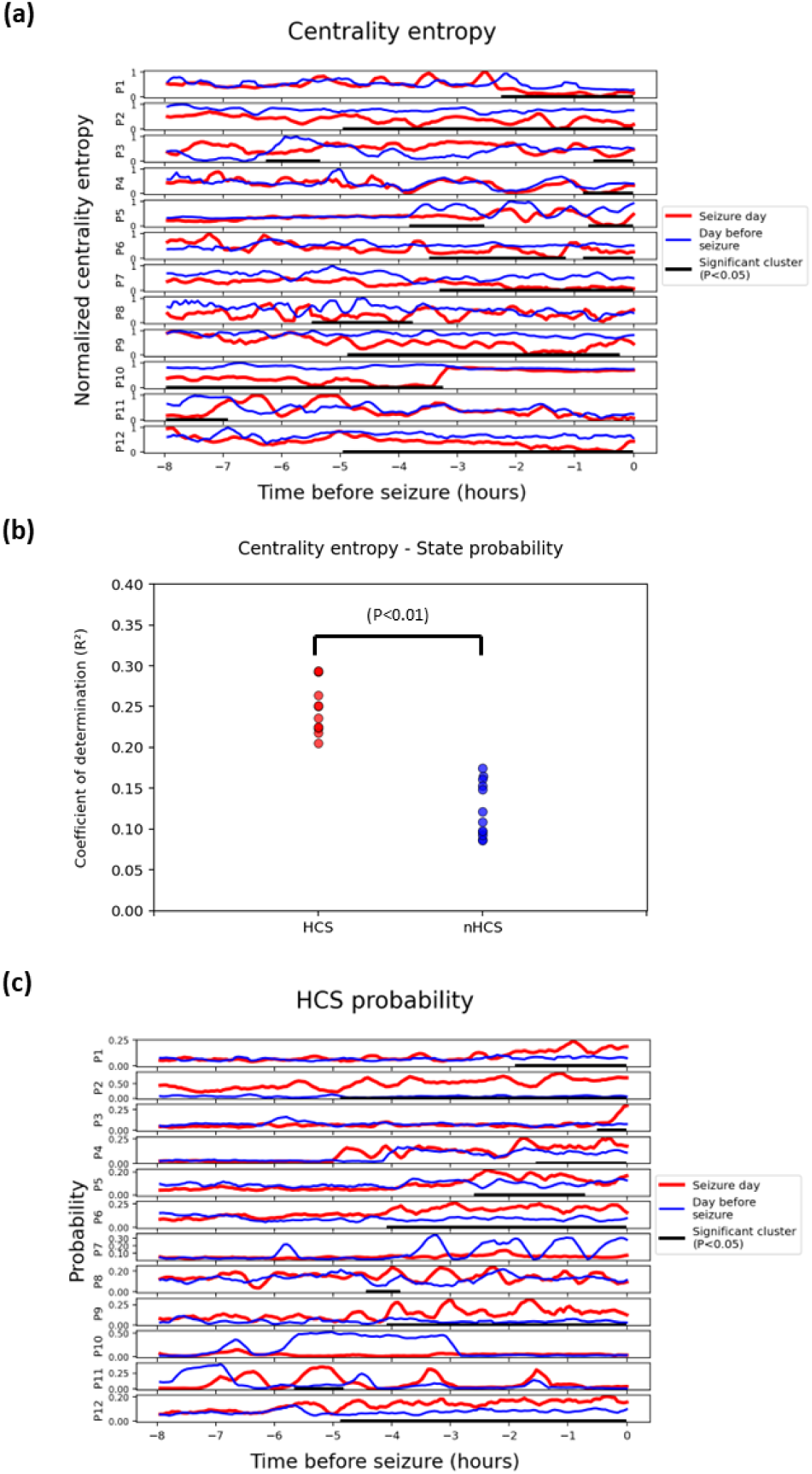
Extension of the HCS analysis to the SWEC-ETHZ iEEG database. This figure shows the dynamics of the centrality entropy, the coefficient of determination and the HCS probability for all patients. In addition to patients P1 and P2, we have added ten patients (P3–P12) from the publicly available SWEC-ETHZ iEEG Database to our analysis, meeting the criterion of having at least 32 hours of data recording before the first seizure, were analyzed. 3(a) The normalized centrality entropy was calculated for both days (red: seizure day; blue: day before the seizure). Overall, in 10 patients (P1-2, P4-7, P9-10 and P12), a significant decrease in entropy was observed hours before seizure onset (Cluster-based permutation test, 𝑃 < 0.05). 3(b) The patient-averaged coefficient of determination (𝑅^2^) of the HCS and nHCS was calculated only for patients of the publicly available database, where each patient may have a different number of network states. It is observed that there is a significant increase (n=10, Mann-Whitney U paired Test, 𝑃 < 0.01) in the HCSs. 3(c) The probability of high-connectivity states (HCS) was calculated for both days. Overall, a significant pre-seizure increase in HCS probability was observed in 10 patients (P1-6, P8-9, and P11-12).

### Robustness across montages

To assess the robustness of our findings to different electrode referencing schemes, we evaluated three standard montages: monopolar (used as baseline), bipolar, and common average reference (CAR). While monopolar references emphasize global effects, bipolar referencing enhances localized activity detection, and CAR mitigates global noise (Ball et al., 2009, Tian et al., 2023, Lachaux et al., 2003).

Analyzing data from Patient 1, Figure S3(a) compares the coefficient of determination values for the three montages in the 8 hours preceding seizure onset. Across all cases, the HCS (State 1) remained the state most strongly associated with centrality entropy changes. Additionally, the time-varying HCS probability analysis (Fig. S3(b)) confirmed a significant pre-seizure increase (∼2 hours before onset) across all montages. In this case, the bipolar montage exhibited the highest 𝑅^2^values and the most pronounced HCS probability increase in P1.

Finally, we validated this analysis in Patient 2. As shown in Fig. S4, the monopolar and common average reference montages demonstrated increased HCS probability, while the bipolar montage did not. This discrepancy may reflect the use of electrocorticography (ECoG) in this patient, as bipolar referencing is less effective for grid electrode topologies due to the inherent spatial configuration of planar electrode arrays (Sazgar et al., 2019).

### Relationship of HCS probability with other signal and network measurements

We evaluated the correlation between HCS probability and other signal-based variables with putative physiological interpretation (mean sEEG signal power at different frequency bands, Figure 4(a); heart rate, Fig. 4(b)) as well as connectivity-based variables (mean sEEG connectivity, Fig. 4(c); mean sEEG centrality, Fig. 4(d)). For each case, we first estimated the coefficient of determination (𝑅^2^) between the probability of HCS and the corresponding variable (mean power, heart rate, etc.) over the time samples accumulated during the 8 hours preceding the seizure onset time on both days (Left panels of Fig. 4; red: seizure day, blue: day before seizure). To examine how the correlation evolved over the above-considered 8 hours, we estimated the time-varying Spearman correlation (𝑟) across windowed estimations of both HCS probability and the corresponding variable (Right panels of Fig. 4; red: seizure day, blue: day before seizure). Specifically, correlation values were estimated for each hour using 3-minute window samples with a step of 15 minutes. In addition, we represented the positive and negative threshold of the obtained the correlation coefficients based on their statistical significance (Fig. 4(b), |𝑟| > 𝑟_𝑡ℎ_ = 0.37, 𝑃 < 0.05).

**Fig. 4.**
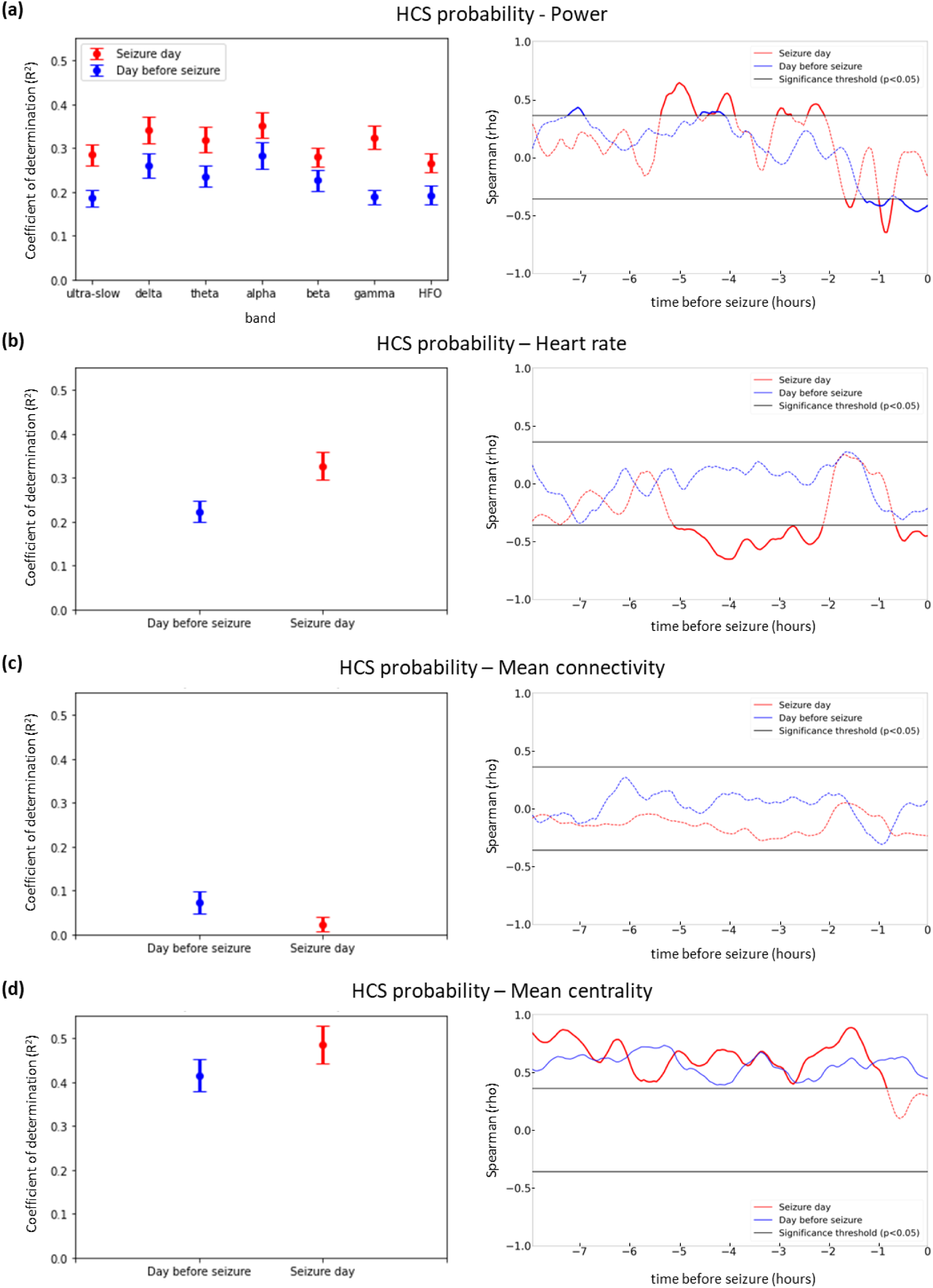
Correlation between HCS probability and signal and connectivity-based variables (P1). The coefficient of determination (left panels) and the time-varying Spearman correlation (Spearman *rho*, right panels) between the probability of HCS and four different measurements on the seizure days (red) and on the day before seizure (blue) are computed. In the left panels, bootstrapping resampling was used to determine a confidence interval for the estimation of the coefficient of determination (𝑅^2^). In the right panels, black lines indicate the significance threshold (𝑃 < 0.05). 4(a) Left: Relationship between HCS probability and normalized power estimation at different physiological bands (ultra-slow, 0.08–0.4 Hz; delta, 1–4 Hz; theta, 4–8 Hz; alpha, 8-12Hz, beta, 12-30Hz; gamma 30–80 Hz; HFO, High-frequency oscillations, 80-150 Hz). Right: Time-varying correlation between the probability of HCS and normalized gamma power. 4(b) Left: Relationship between HCS probability and the heart rate. Right: Time-varying correlation between the probability of HCS and the heart rate. 4(c) Left: Relationship between HCS probability and the mean connectivity. Right: Time-varying correlation between the probability of HCS and the mean connectivity. 4(d) Left: Relationship between HCS probability and the mean centrality. Right: Time-varying correlation between the probability of HCS and the mean centrality.

The analysis of mean sEEG power revealed that HCS probability exhibited the strongest association with the ultra-slow (0.08–0.4 Hz), delta (1–4 Hz), theta (4–8 Hz), and gamma (30– 80 Hz) bands (Fig. 4(a) left). The gamma band was further analyzed using time-varying Spearman correlation, which showed significant discontinuous positive correlations with HCS probability hours before seizure onset. However, near the seizure onset, this relationship transitioned to a negative correlation (Fig. 4(a) right). In the case of heart rate, the overall association with HCS probability was moderate, with 𝑅^2^ values below 0.4 (Fig. 4(b) left). In contrast, the time-varying analysis revealed two distinct periods of significant negative correlation between heart rate and HCS probability: a sustained period from 5 to 2 hours before seizure onset and a brief period immediately preceding seizure onset. These findings suggest that increases in HCS probability during these periods coincide with reductions in the patient’s heart rate (Fig. 4(b) right). Additionally, we replicated this analysis in Patient 2, observing a comparable increase in Spearman correlation on seizure days (Fig. S6).

Mean functional connectivity showed very low 𝑅^2^ values and no significant time-varying correlation, indicating that the dynamics of HCS, as inferred over short timescales (600 ms), are largely independent of the grand-average sEEG connectivity (Fig. 4(c)). Finally, the analysis of mean centrality revealed relatively high 𝑅^2^ values on both days, reflecting that HCS are likely to represent a class of eigenvectors with large mean centrality. Accordingly, this association remained largely unspecific to the seizure day (Fig. 4(d)).

In summary, the strongest and most genuine association with HCS probability was observed for heart rate, suggesting a possible link between network connectivity and autonomic function. This relationship should be further investigated in larger patient cohorts, as it may reveal novel non-invasive biomarkers for pre-seizure periods. In particular, understanding the mechanistic basis for this association, particularly the role of cerebral blood flow and its connection to network synchronizations, could provide valuable insights into seizure dynamics.

### Pre-seizure HCS probability is driven by changes in SOZ network centrality

In Patient 1, the pre-seizure occurrence of more frequent high-connectivity states (HCS) (see Fig. 5(a) for examples of HCS and non-HCS samples) starting two hours before seizure onset (Fig. 5(b), top) was not fully associated with a consistent increase in the time-varying mean connectivity (Fig. 5(b), middle). This observation aligns with our earlier findings (Tauste Campo et al., 2018) and the correlation analysis presented in Fig. 4(c). In contrast, the time-varying mean centrality showed a pronounced deviation relative to the corresponding period on the previous day (Fig. 5(b), bottom). This prompted us to examine how the dynamics of node centrality influenced the emergence of HCS in this patient. To address this question, we first analyzed the mean centrality distribution across different types of channels, distinguishing between channels within the seizure-onset zone (SOZ), as identified by two epileptologists (AD, MK), and the remaining channels. Notably, the mean centrality of SOZ channels displayed a substantial increase during the two hours preceding the seizure compared to the previous day, mirroring the pattern of the overall mean centrality. In contrast, the mean centrality of non-SOZ channels exhibited more subtle alterations with respect to the previous non-seizure day (Fig. 5(c), top). A similar effect was also observed in Patient 2 (P2) four hours prior to seizure onset (Fig. 6(b), bottom), further supporting our hypothesis.

**Fig. 5.**
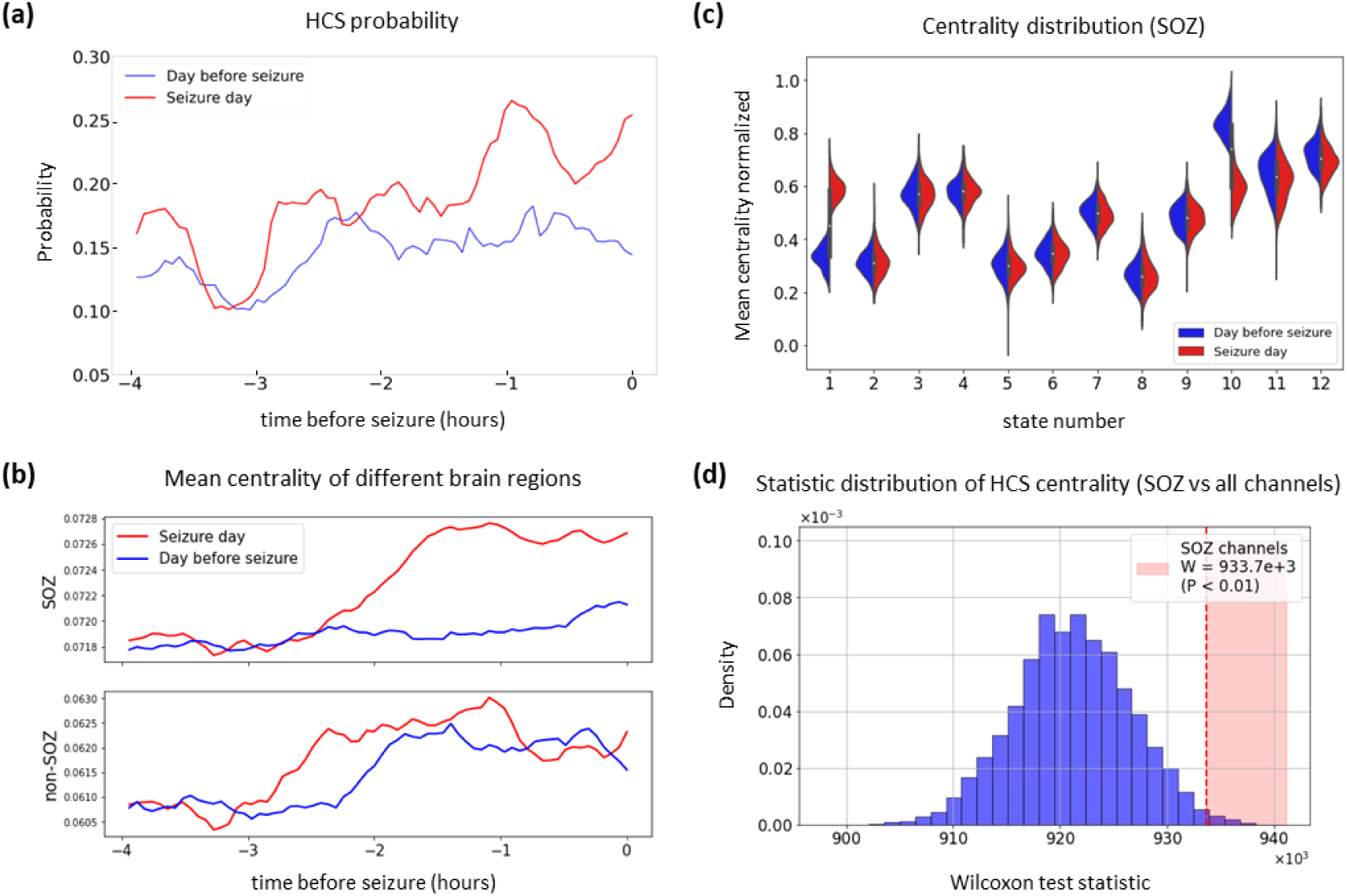
Influence of the SOZ dynamics on pre-seizure HCS probability (P1). 5(a) Time-varying HCS probability over the 4 hours preceding seizure onset on the seizure day (red) and the day before (blue). 5(b) Time-varying mean centrality for SOZ channels (above, 𝑛 = 11) and random non-SOZ (below, 𝑛 = 11) channels. 5(c) Distribution of mean centrality for the SOZ across the 12 identified network states. 5(d) Surrogate distribution of the Wilcoxon test statistic built using 11 randomly selected non-SOZ channels, with the original test statistic highlighted by a red vertical dashed line (𝑃 < 0.01).

**Fig. 6.**
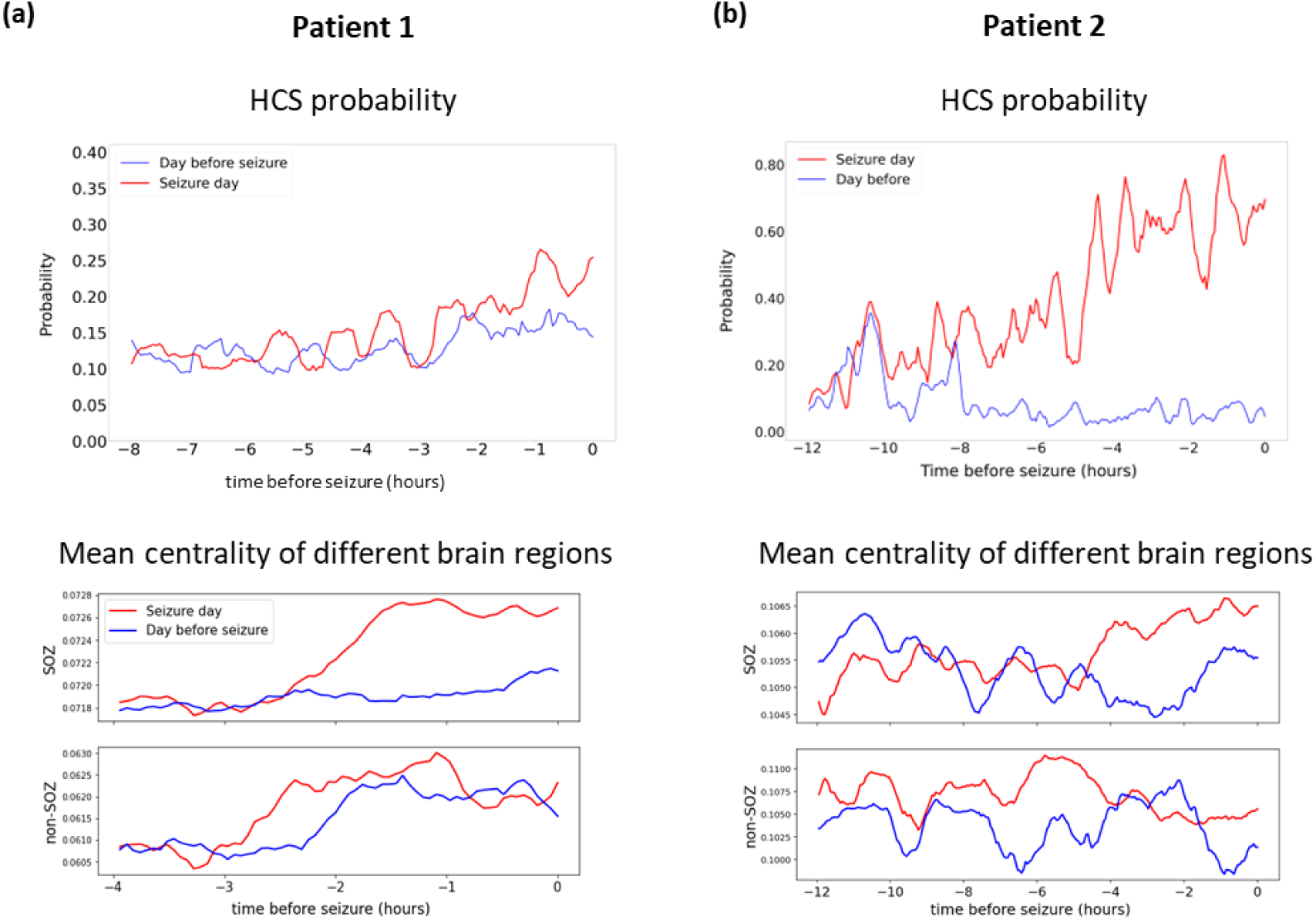
Influence of the SOZ dynamics on pre-seizure HCS probability (Comparison between P1 and P2). 6(a) The HCS probability and the mean centrality of each brain region are shown. 6(b) Top: In P2 it is observed that the HCS probability increases as the seizure approaches, specifically, this happens from 8 hours before the onset of the seizure. 6(b) Bottom: it shows an increase in the mean centrality located in the SOZ channels that is evident from 5 hours before the onset of the seizure.

Next, we investigated whether the between-day increase in SOZ mean centrality was specifically associated with the HCS. An analysis across the 12 original states revealed that the pre-seizure increase in SOZ centrality was exclusively tied to the HCS (Fig. 5(c), middle). To confirm that this specificity was reciprocal, we compared the centrality of SOZ channels to a random subsample of channels of equal size. The increase in SOZ centrality was statistically significant (P < 0.01) (Fig. 5(c), bottom). These results, also replicated in Patient 2 (see Fig. S7), suggest that the emergence of HCS during the pre-seizure period is specifically driven by an increase in the mean centrality of SOZ channels.

At first glance, it may seem counterintuitive that states with identical mean connectivity can exhibit different mean centrality values. This discrepancy can be explained by a network model in which rearrangements within the connectivity matrix alter the centrality of certain nodes while preserving the overall connectivity (See next Section). For instance, in Patient 1, our data analysis revealed that no significant changes in mean connectivity were observed across days for every state, and only the centrality of SOZ channels in HCS was increased during the pre-seizure period (Fig. S5).

In conclusion, our findings emphasize the critical role of SOZ-specific channels in driving the dynamics of HCS probability. Assessing the topological dynamics of these channels offers deeper insights into network-level changes that precede seizure onset and highlights the SOZ as a key contributor to the emergence of pre-seizure HCS.

### A low-dimensional stochastic network explains how topological reconfigurations increase pre-seizure HCS probability

To further interpret the relationship between the dynamics of SOZ centrality and HCS probability during the pre-seizure period, we proposed a low-dimensional stochastic network model (N = 4) designed to simulate the key network dynamics illustrated in Fig. 5. This model generated sequences of covariance matrices within a controlled range of mean connectivity values ensuring time continuous signals within each day and similar connectivity distributions across both days (Fig. 7(b), top right). However, during the two hours preceding seizure onset, the centrality distribution was altered between the two days by strategically rearranging the positions of correlation coefficients within the covariance matrices (Fig. 7(a)). Specifically, to manipulate mean centrality while maintaining constant mean connectivity, the coefficients within the correlation matrix were reorganized. This reconfiguration allows two matrices with identical mean connectivity to be classified into distinct states: high-connectivity states (HCS) and non-HCS. This distinction arises because the classification of network states prioritizes the structure of centrality (via eigenvectors) before ranking based on mean connectivity values. Fig. 7(a) demonstrates this principle: reorganizing the correlation coefficients to emphasize a hub channel (e.g., Ch1) increases the mean centrality (as observed on the seizure day), while an even distribution of coefficients produces a lower mean centrality (as seen on the day prior to seizure onset).

**Fig. 7.**
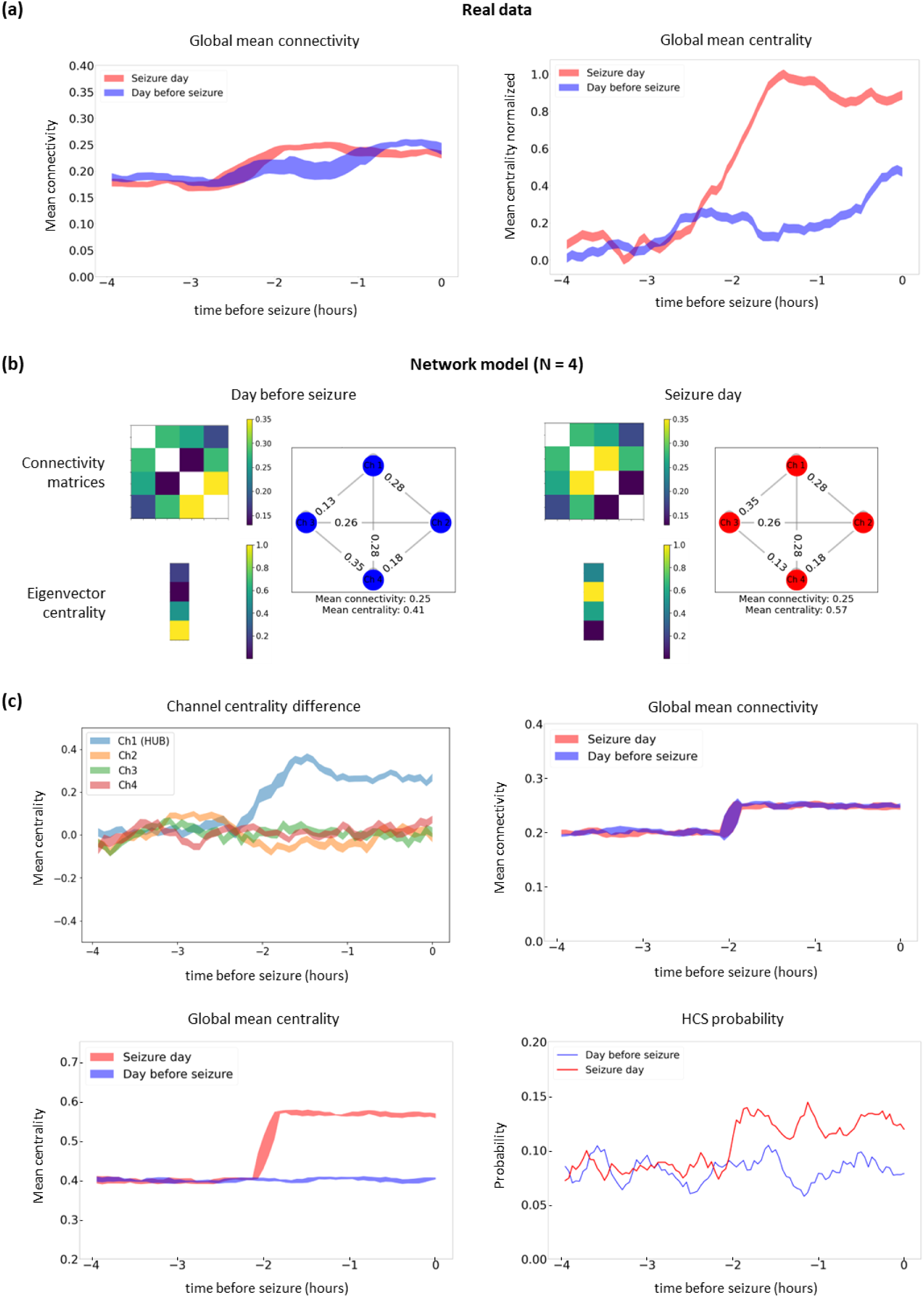
Low-dimensional stochastic network model reproducing the preictal emergence of HCS probability. (a) The global mean connectivity and centrality of patient P1 are presented, illustrating how dynamic shifts in centrality—rather than connectivity—guided the development of our stochastic network model. (b) The model generates random correlation matrices (𝑁 = 4) with fixed mean connectivity and varying mean centrality by rearranging correlation coefficients. This rearrangement produces two network states: one with uniform coefficient distribution (Left panel: lower centrality, as observed the day before the seizure) and another with a hub-like structure (Right panel: higher centrality, resembling the seizure day). Network states are classified based on eigenvector centrality, prioritizing centrality over connectivity. (c) Results averaged over 40 simulation runs. The four subfigures are interrelated, as they arise from the same set of simulations. The top left figure shows that HCS probability (bottom left) increases significantly only when a specific channel (e.g., Ch1) exhibits dominant centrality, forming a hub (analogous to the clinical SOZ). This centrality dominance explains the observed rise in HCS probability during the 2 hours preceding a seizure. The corresponding global mean connectivity and mean centrality are presented in the top right and bottom right figures, respectively. Together, these results demonstrate that the increase in HCS probability emerges as a topological property of the model.

To explore the effects of these centrality changes on HCS probability, we analyzed the centrality dynamics of individual channels (Fig. 7(b) top left). When no single channel displayed prominent centrality (e.g., two to four hours before the seizure), the HCS probability remained relatively stable (Fig. 7(b), bottom right). In contrast, when a specific channel (Ch1) dominated centrality, a significant increase in HCS probability was observed during the two hours leading up to the seizure (Fig. 7(b), bottom right). These results suggest that, within the model, channels exhibiting high centrality, analogous to the clinical seizure-onset zone (SOZ), play a key role in modulating both global mean centrality and HCS probability. This behavior mirrors the real data, where the clinical SOZ channels predominantly shape the network’s mean centrality dynamics and drive the emergence of HCS. The global mean connectivity and centrality remained stable over the simulated period (Fig. 7(b), top right and bottom left, respectively), reinforcing the interpretation that centrality changes—rather than connectivity variations—are responsible for pre-seizure increases in HCS probability.

The strength of this low-dimensional stochastic network model lies in its ability to capture the observed pre-seizure behavior as an intrinsic network property. Specifically, the model demonstrates that an increase in HCS probability already arises when overall centrality becomes concentrated in a single channel as seizure onset approaches. This finding is further validated by averaging results across 40 independent simulations of the model, which replicates the pre-seizure emergence of HCS seen in the experimental data.

## 4. DISCUSSION

In this study, we investigated the occurrence of high-connectivity states (HCS) as a potential network-level biomarker of pre-seizure dynamics using intracranial EEG recordings from 12 patients with drug-resistant epilepsy. Building on previous research (Tauste Campo et al., 2018), we validated and expanded those findings in a larger and more diverse patient cohort. Our results revealed a significant pre-seizure decrease in centrality entropy driven by increased HCS probability. In two patients with validated seizure-onset zones (P1 and P2), we studied the association between the time-varying HCS probability and the network relevance of SOZ with respect to the remaining sites. These findings across both patients motivated the development of a low-dimensional stochastic network model to simulate the observed common pre-seizure dynamics. Additionally, we examined the robustness of HCS probability under different referencing montages and analyzed its correlation with other signal and connectivity-based variables including the heart rate.

### Replication and generalization

We reproduced our earlier findings across 12 patients from three different sources, including two patients (P1 and P2) with clinical and additional psychological data: patient P1 from a collaborating hospital, patient P2 from IEEG.ORG database and patients P3–P12 from SWEC-ETHZ iEEG long-term database. In 9 out of 12 patients, centrality entropy significantly decreased within up to 8 hours preceding seizure onset, a reduction that we attributed to an increase in HCS probability. However, the 10 patients from the online database provided limited metadata, including seizure onset times but lacking crucial clinical details such as electrode labels, co-registered electrocardiogram (ECG), and validated seizure-onset zones. As a result, a more detailed analysis was performed only in P1 and P2, where richer clinical and recording data were available. For these patients, we explored the robustness of HCS probability across different referencing montages, investigated its correlation with signal and connectivity-based variables, and assessed the influence of SOZ dynamics on HCS probability.

### Robustness across montages and correlation with other signal-based variables

The robustness of HCS probability was confirmed across monopolar, common average, and bipolar sEEG montages, where we consistently observed significant increases in HCS probability hours before seizure onset, with high coefficients of determination. To further interpret HCS probability within a broader physiological and functional connectivity framework, we compared it with other previously studied variables, including sEEG signal power (Vila-Vidal et al., 2017), heart rate (Karoly et al., 2021), mean connectivity, and mean centrality. While HCS probability increased almost uniformly across physiological frequency bands, gamma-band power correlations fluctuated near significance thresholds over time. In contrast, heart rate displayed sustained significant correlations between 2 to 5 hours and again 30 minutes before seizure onset. Mean connectivity consistently exhibited low correlations with HCS probability, while mean centrality showed relatively higher correlations. These results suggest that heart rate and mean centrality could complement HCS probability, providing a multidimensional perspective for predicting epileptic seizures (Schlegel et al., 2024).

### Topological reconfiguration drives the emergence of high-connectivity states: insights from a low-dimensional stochastic network model

To better understand the topological mechanisms underlying the emergence of high-connectivity states (HCS), we developed a low-dimensional stochastic network model (𝑁 = 4 nodes) that captures key topological changes in brain connectivity. Network dynamic models are particularly valuable for analyzing the time-varying interactions that drive seizure generation (Sriraam et al., 2018). In our study, we observed that both mean connectivity and mean centrality increased two hours prior to seizure onset; however, mean centrality showed a larger change with respect to the previous non-seizure day (e.g., in P1). This observation led to the design of a network model based on covariance matrices that preserved mean connectivity but allowed variations in centrality by rearranging the positions of correlation coefficients.

The model demonstrated that changes in mean centrality, rather than mean connectivity, might drive the emergence of HCS probability. Specifically, when centrality was evenly distributed across the network, HCS probability remained stable during the pre-seizure period (2–4 hours before seizure onset). However, when centrality became concentrated in a single hub node, HCS probability significantly increased closer to seizure onset (0–2 hours before the event). Averaging across a number of simulations, we found that enhanced HCS probability emerged as an intrinsic property of the network when centrality dynamics became dominated by a subset of nodes, resembling the behavior of SOZ channels observed in real data. Thus, the network model highlights the critical role of centrality dynamics in driving pre-seizure network behavior.

### Study limitations

Despite the promising findings of this study, several limitations must be acknowledged. While heart rate and mean centrality emerged as potential complementary biomarkers to HCS probability, these observations were derived from two patients (P1 and P2) with detailed clinical and physiological data. This limitation prevented us from integrating these variables into a unified predictive model. Future studies should address this by including a larger cohort of patients with more complete metadata, such as epilepsy type, intracranial EEG montages, medication effects, and data from multiple seizures beyond the first monitored event. Another limitation lies in the use of Pearson correlation to measure functional connectivity, which is effective for capturing linear relationships but may fail to identify nonlinear signal dependencies. Exploring alternative connectivity measures that account for nonlinear interactions, such as phase-based or mutual information approaches, could reveal further insights into pre-seizure dynamics (Mercier et al., 2024).

The role of spontaneous interictal spikes, which are closely associated with seizure onset, also remains an important consideration for future research. Examining their distribution and behavior over extended pre-seizure periods may uncover additional predictive biomarkers for seizure onset (Karoly et al., 2016). Furthermore, heart rate variability (HRV) has been identified as a reliable pathological indicator of epilepsy (K. Li et al., 2023), and its integration into predictive frameworks should be explored. Lastly, the use of our low-dimensional stochastic network model provides an important avenue for simplifying network-level analyses. Future work should resort to such models to determine the minimum number of channels necessary for accurate seizure prediction. Reducing the dimensionality of seizure prediction models could improve clinical feasibility and computational efficiency, particularly in real-world applications (Statsenko et al., 2023).

## 5. CONCLUSION

In conclusion, our findings provide evidence that time-dependent increases in HCS probability throughout the day can serve as a potential pre-seizure biomarker. By combining detailed sEEG and ECoG analysis with a low-dimensional network model, we propose that the preictal emergence of HCS reflects an intrinsic property of network organization primarily governed by centrality. Future research efforts aimed at integrating complementary physiological and connectivity-based measures into a unified, multivariate predictive framework could further enhance the clinical utility of HCS probability for reliable early detection of upcoming seizures.

## Acknowledgements

N.M., M.V-V. and A.T.C were supported by the National Research project PID2020-119072RA-I00/AEI/10.13039/501100011033) funded by the Spanish Ministry of Science, Innovation, and Universities. N.M was supported by the predoctoral grant program “FI-SDUR” from the Department of Research and Universities, Catalan Government, 2022 (ref. BDNS 612831) (DOGC Núm. 8621 - 8.3.2022). M.V-V. was supported by the grant PTQ2022-012679, funded by the Spanish Ministry of Science, Innovation, and Universities.

## Ethical statement

All diagnostic and surgical procedures were approved by The Clinical Ethical Committee of Hospital Clínic and all clinical investigation was conducted according to the principles expressed in the Declaration of Helsinki. Following the Declaration of Helsinki, the patient was informed about the procedure, and they gave their written consent beforehand.

## Supplementary figures

**Fig. S1.**
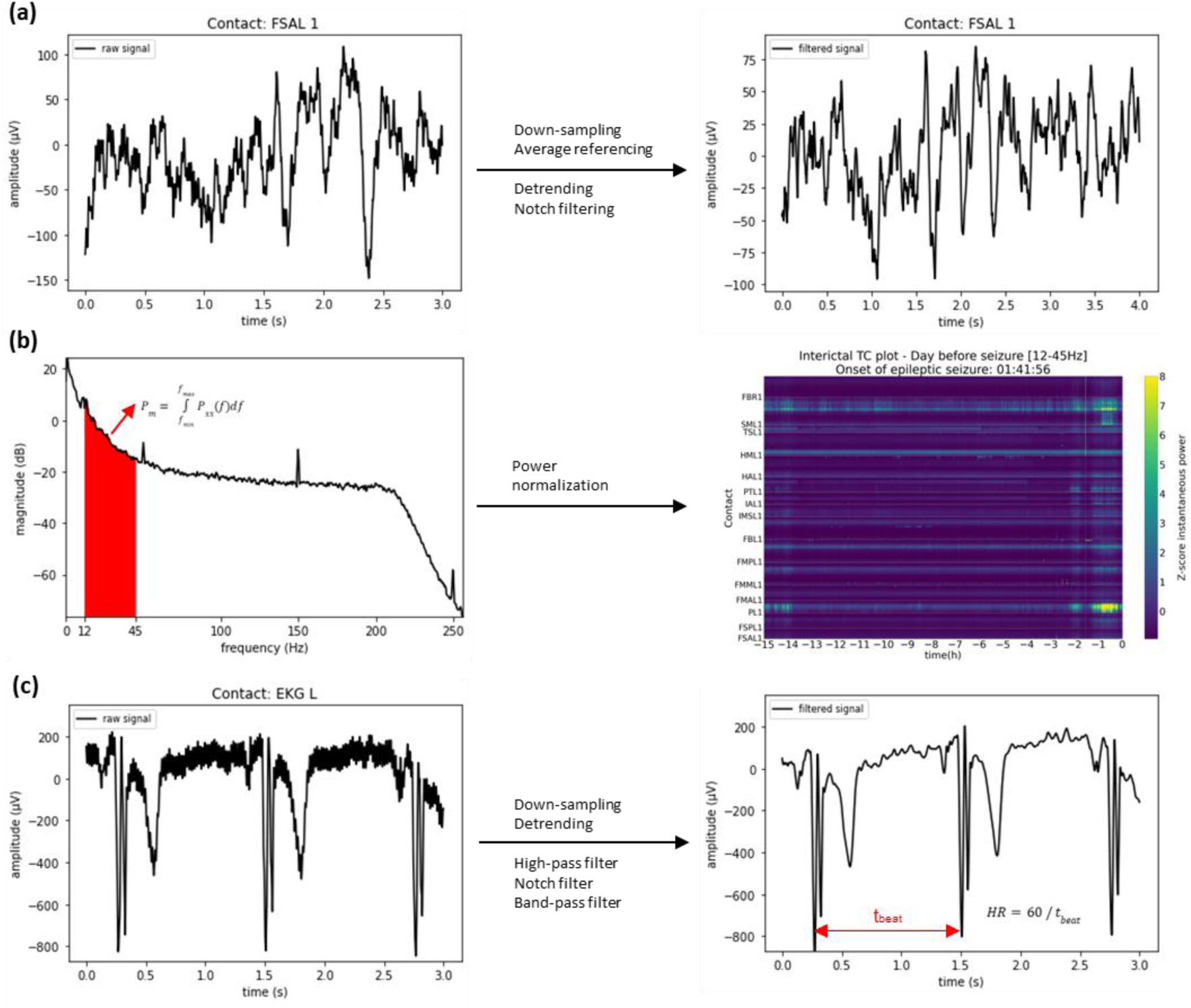
Power and heart rate analysis. S1(a) sEEG signal preprocessing. Left: An exemplary raw sEEG signal is represented. Right: Preprocessing techniques (down-sampling, average referencing, detrending, and notch filtering) are applied to obtain a cleaner sEEG signal for analysis. S1(b) Power analysis. Left: The power spectral density (PSD) of the filtered signal is calculated and then integrated over the frequency range of interest (e.g. 12-45 Hz) to calculate power. Right: The power is normalized by computing its Z-score relative to a baseline defined as the first 5 minutes of the initial recording day. This spatio-temporal representation (time on the 𝑥-axis and channel contacts on the 𝑦-axis) illustrates an increase in power in specific channels (yellow) as the seizure approaches. S1(c) Heart rate analysis. Left: An exemplary raw ECG signal is represented. Right: Preprocessing steps (down-sampling, high-pass filtering [<0.5 Hz], notch filtering, band-pass filtering [0.5-40 Hz]) are applied to detect the time between 𝑅 -peaks (𝑡_𝑏𝑒𝑎𝑡_) and calculate heart rate based on 𝑡_𝑏𝑒𝑎𝑡_.

**Fig. S2.**
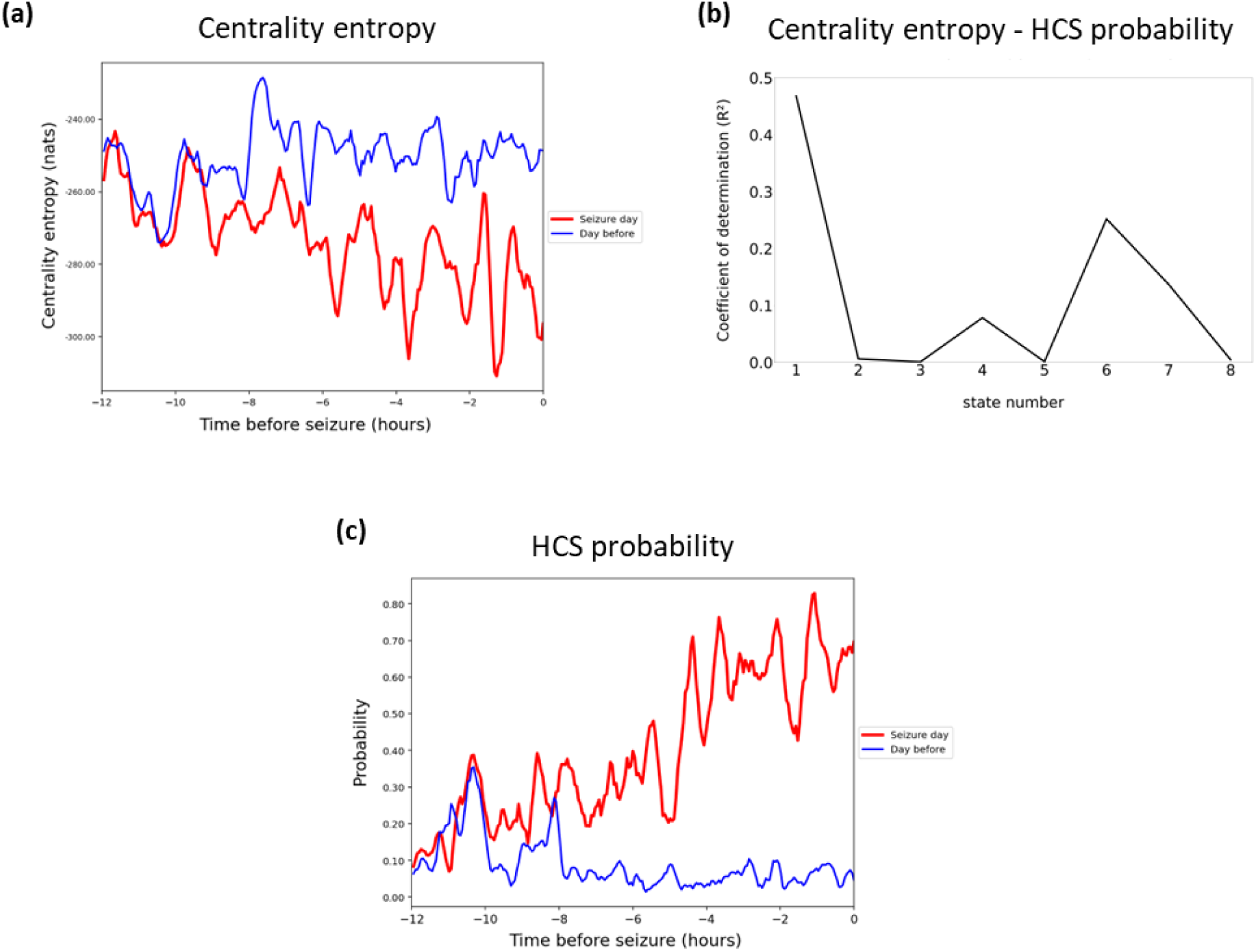
Increased probability of high connectivity states (HCS) explains changes in pre-seizure ECoG network dynamics variability (Patient 2). The results illustrated in Fig. 2a), 2c) and 2e) are reproduced for Patient 2 in Fig S2 a), b) and c), respectively.

**Fig. S3.**
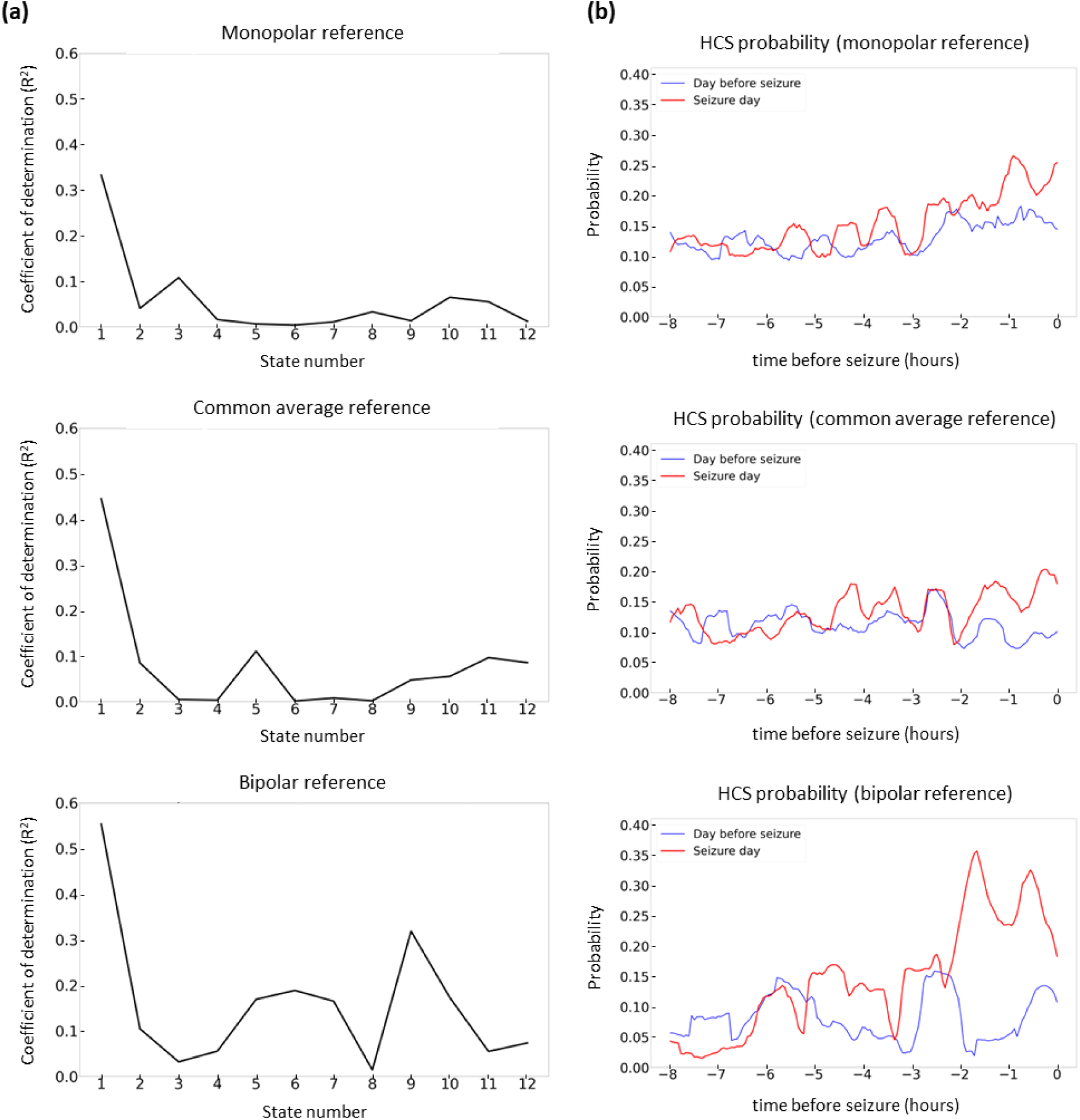
Robustness of HCS pre-seizure emergence to reference montages (Patient 1). Evaluation of HCS probability across three different reference montages: monopolar (top panels), common average (middle panels), and bipolar (bottom panels). The patient data were initially recorded with a monopolar reference. For the common average reference, the mean of all channels was subtracted from each individual channel, while for the bipolar reference, consecutive channels from the same electrode were subtracted. S3(a) The coefficients of determination (𝑅^2^) between centrality entropy and the probability of each state were calculated for each reference. In all montages, state 1 (HCS) best explained the entropy drop observed hours before seizure onset. The results remained consistent across reference configurations, with the highest R² value obtained in the bipolar montage (bottom panel). S3(b) The dynamics of HCS probability over the 8 hours prior to seizure onset for both the seizure day and the previous day are shown for each reference montage. In all cases, an increase in HCS probability as compared to the previous day was observed in the 2 hours leading up to seizure onset.

**Fig. S4.**
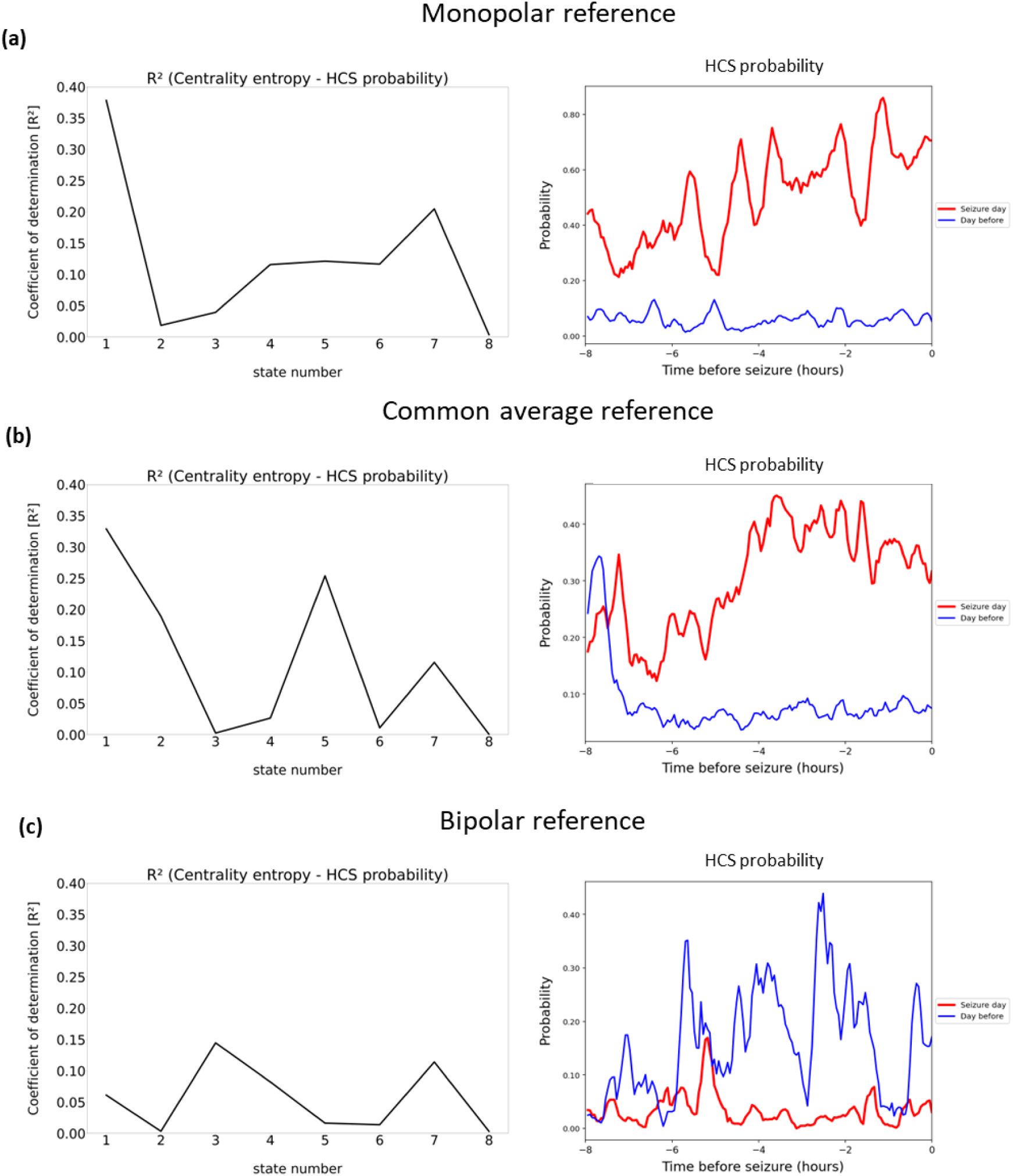
Robustness of HCS pre-seizure emergence to reference montages (Patient 2). This figure presents a replication of the analysis shown in Fig. S3, demonstrating that the HCS probability increases as the seizure approaches when using either monopolar or common average reference schemes. This preictal pattern was not observed in the bipolar reference montage. We hypothesize that this discrepancy may reflect fundamental differences in signal acquisition between electrocorticography (ECoG) and stereoelectroencephalography (sEEG), as the implanted electrodes in this patient were ECoG grids rather than sEEG depth electrodes.

**Fig. S5.**
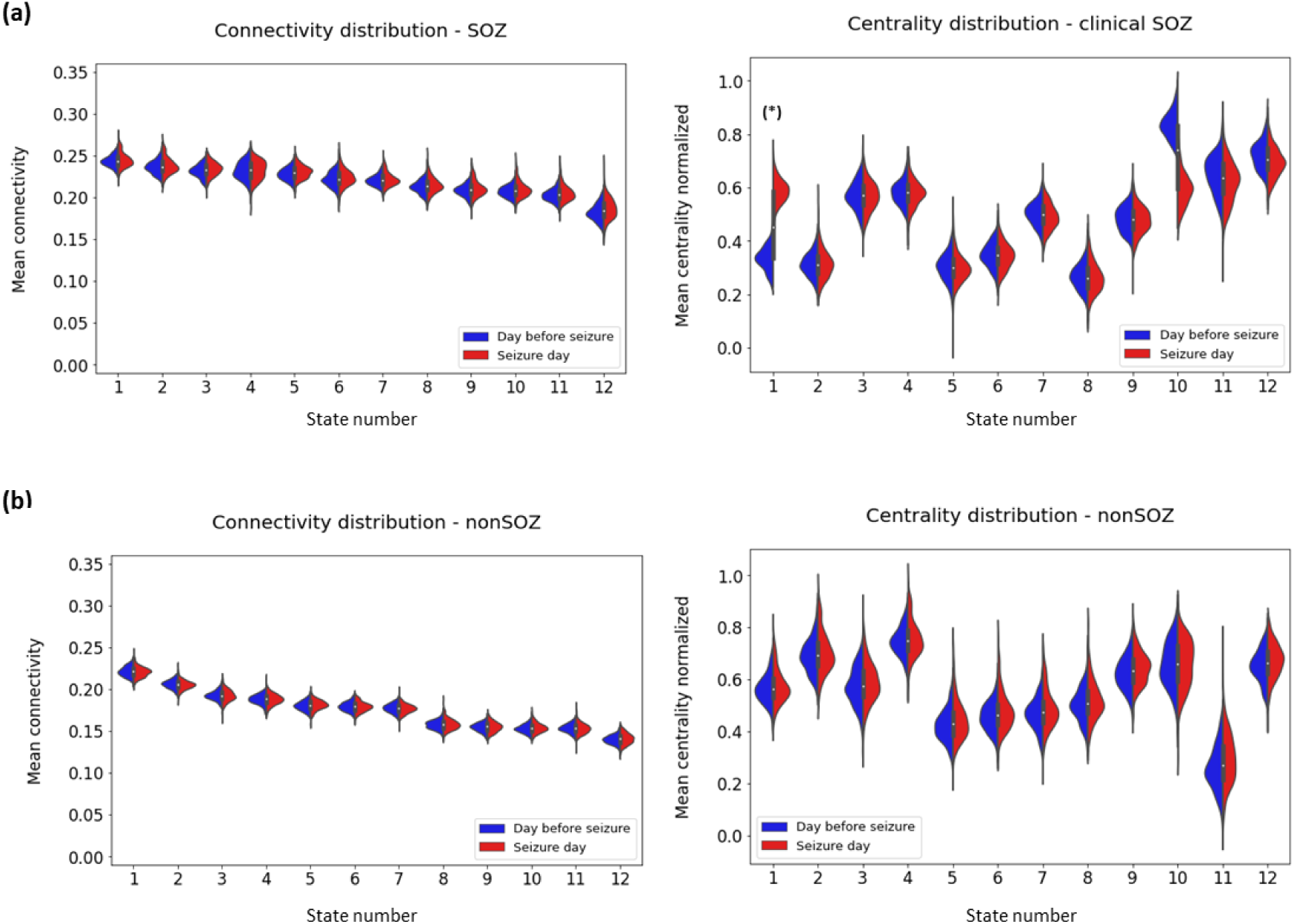
Comparison of mean connectivity and mean centrality for SOZ and non-SOZ regions (P1). The distributions of mean connectivity and mean centrality for the SOZ (A) and non-SOZ (B) channels during the 8 hours preceding the seizure are represented, with the seizure day shown in red and the previous day in blue. S2(a) SOZ channels. Left: The mean connectivity did not change significantly across both days for any network state. Right: A significant increase in centrality was observed in state 1, corresponding to the HCS (one-sided Wilcoxon test, 𝑃 < 0.01). Additionally, a difference of opposite signs arose in state 10 on the seizure day. S2(b) non-SOZ channels. Left: No significant changes in mean connectivity were observed. Right: No significant changes in mean centrality were observed.

**Fig. S6.**
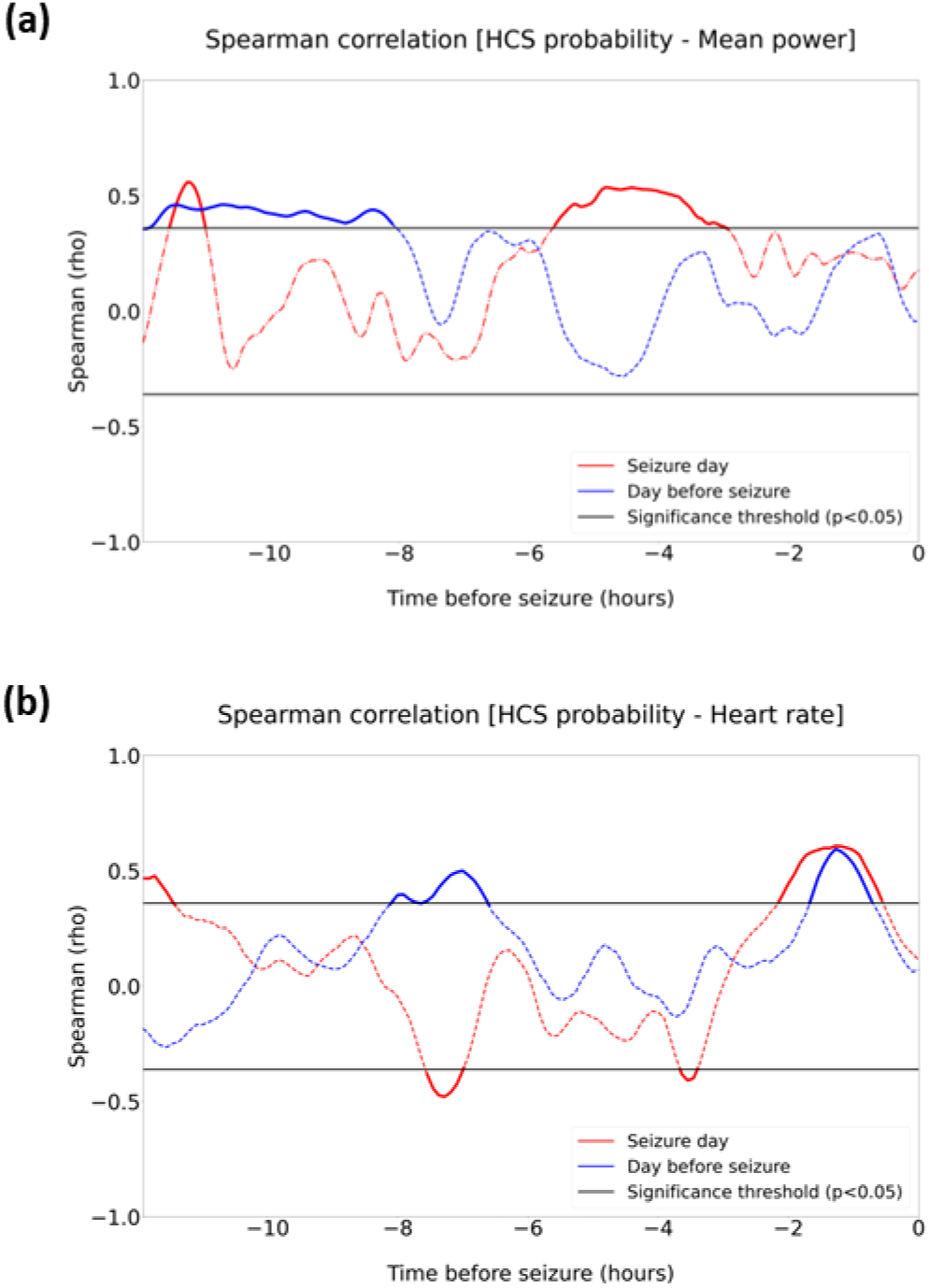
Time-varying correlation of HCS probability with gamma power oscillations and heart rate fluctuations (Patient 2). Time-varying Spearman correlation (Spearman *rho*) between the probability of HCS and two signal-based measurements on the seizure days (red) and on the day before seizure (blue) are computed over up to 12 hours preceding seizure-onset time. (a) Spearman correlation between HCS probability and mean gamma-band power (b) Spearman correlation between HCS probability and the heart rate.

**Fig. S7.**
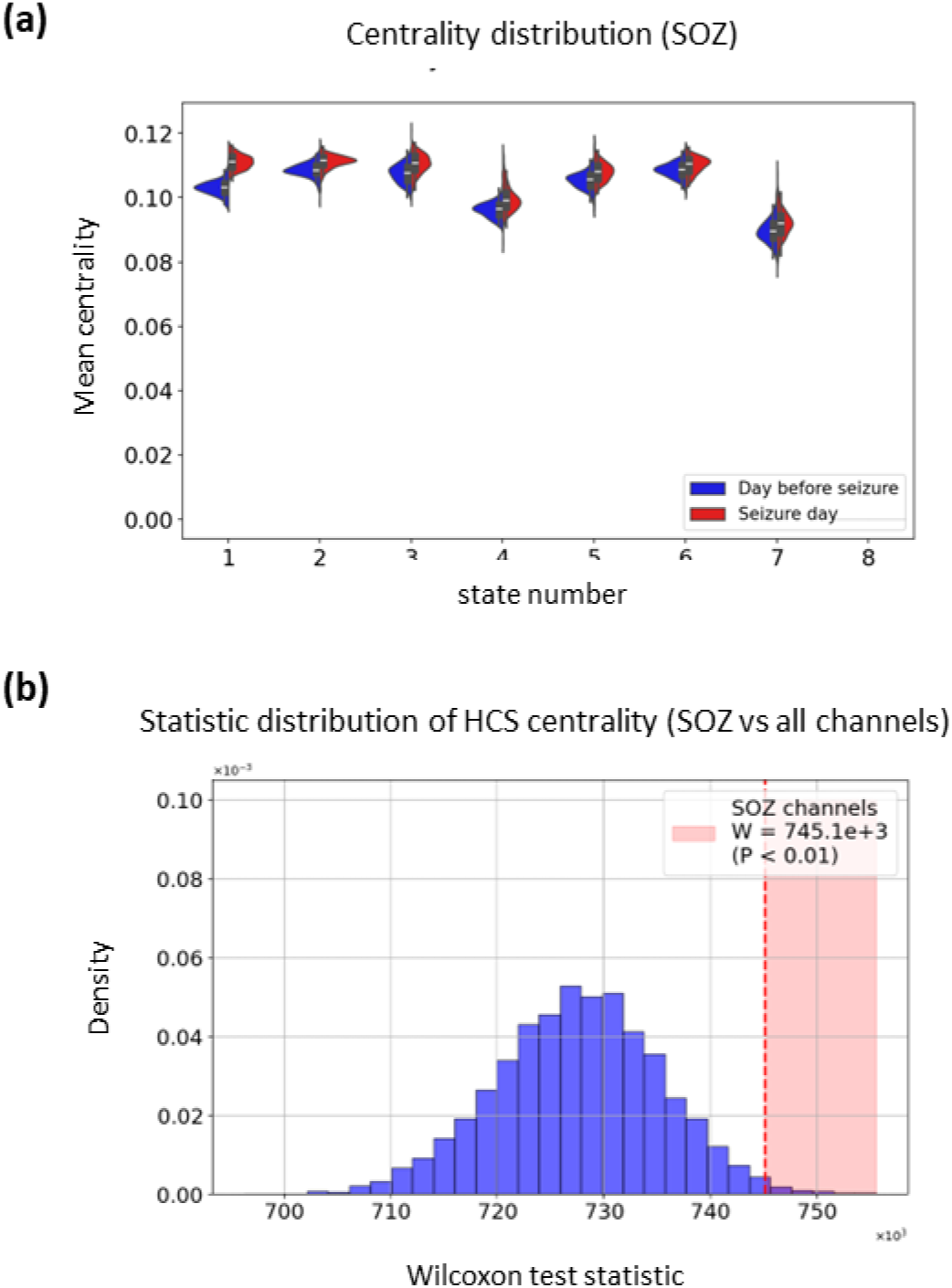
Specific association between SOZ and HCS (Patient 2). (a) Distribution of mean centrality across the 7 identified network states. (b) Surrogate distribution of the Wilcoxon test statistic built using 10 randomly selected non-SOZ channels, with the original test statistic highlighted by a red vertical dashed line (𝑃 < 0.01).

